# *SMPD3* suppresses IDH mutant tumor growth via dual autocrine-paracrine roles

**DOI:** 10.1101/2020.07.14.202200

**Authors:** Anjali Balakrishnan, Fermisk Saleh, Lata Adnani, Vorapin Chinchalongporn, Ahmed El-Sehemy, Thomas Olender, Myra J Chen, Shiekh Tanveer Ahmad, Oleksandr Prokopchuk, Lakshmy Vasan, Yacine Touahri, Rehnuma Islam, Sajeevan Sujanthan, Dawn Zinyk, Lacrimioara C Comanita, Boris Kan, Taylor Fleming, Iacovos P Michael, Cindi M Morshead, Hon S Leong, Satoshi Okawa, Marjorie Brand, Valerie A Wallace, Jennifer A Chan, Carol Schuurmans

## Abstract

Isocitrate dehydrogenase (IDH) mutant gliomas, including oligodendroglioma (IDH-O) and astrocytoma (IDH-A), have signature slow-growth rates that are poorly understood. Here, we reveal that *SMPD3,* a ceramide-producing sphingomyelinase implicated as a tumor suppressor gene and involved in extracellular vesicle biogenesis, suppresses IDH-mutant tumor growth via autocrine and paracrine actions. In patients with IDH-mutant gliomas, higher *SMPD3* expression levels correlate with longer survival, consistent with ceramide acting as an anti-oncometabolite. *SMPD3* knock-down in patient-derived IDH-O cells enhances proliferation cell-autonomously in 2D-culture and 3D-human cerebral organoids, and accelerates tumor growth in mouse orthotopic xenografts. Supporting paracrine actions, IDH-O-derived extracellular vesicles, enriched in ribosomal proteins, induce astrocytic death *in vitro*. Furthermore, non-neoplastic glia in IDH-O tumors proliferate abnormally yet undergo apoptosis, concomitant with the acquisition of a translation-enriched transcriptional signature by tumor-associated oligodendrocytes. *SMPD3* thus suppresses IDH-mutant glioma growth cell-autonomously and phenotypically alters normal glia via extracellular vesicle biogenesis and paracrine actions.

## INTRODUCTION

IDH-mutant gliomas are diffusely infiltrating glial brain tumors that include oligodendrogliomas and astrocytomas^1-4^. Aside from bearing an IDH1 or IDH2 mutation, oligodendrogliomas (IDH-O) are further distinguished genetically by 1p/19q chromosomal arm co-deletions, *TERT* promoter mutations, and CIC mutations ^5^, whereas IDH-mutant astrocytomas (IDH-A), typically carry *TP53* and *ATRX* mutations ^6^. Compared to their IDH-wildtype counterpart glioblastoma multiforme (GBM), IDH-mutant gliomas have more indolent growth and longer median survival times ^7,8^. Why gliomas with IDH mutation have slower growth and less aggressive clinical courses, however, is not well understood ^9^.

One factor implicated in distinguishing IDH-wild-type versus IDH-mutant glioma growth rates is the “sphingolipid rheostat”, which controls the balance between sphingosine-1-phosphate (S1P), a pro-proliferative lipid second messenger, and ceramide, which suppresses cell growth and induces apoptosis ^10-12^. Ceramide acts cell-autonomously to suppress cell proliferation and induce cell death through autocrine signaling and is also secreted, impacting surrounding cells via paracrine signaling. *SMPD3*, which encodes for neutral sphingomyelinase 2 (nSMase2), regulates the sphingolipid rheostat via hydrolyzing S1P to produce ceramide^10^. Therefore, *SMPD3* has been designated as a putative tumor suppressor gene, and its derivative ceramide as an anti-oncometabolite ^13-15^. Among brain cancers, S1P levels are elevated in IDH-wildtype GBM cells, whereas IDH-mutant glioma cells have higher levels of ceramide. Furthermore, in experimental models such as the rat C6 glioma cell line, expression of *Smpd3* inhibits tumour growth ^16^. These data suggest that the sphingolipid rheostat is shifted in opposite directions in IDH-mutant versus IDH-wildtype gliomas – with the balance towards pro-proliferative in GBM and anti-proliferative/pro-apoptotic in IDH-O and IDH-A ^10-12^.

Paracrine signaling occurs by various modes of intercellular communication, including but not limited to, extracellular vesicles (EVs) ^17,18^. Ceramide produced by nSMase2 is involved in the production of EVs ^19,20^, which are secreted by most cells in the brain, including oligodendrocytes, astrocytes, neurons, and cancer cells ^21-24^. EVs package lipids, proteins, and nucleic acids (DNA, mRNA, microRNA, non-coding RNA) in a lipid bilayer that protects cargo from degradation in the extracellular space and facilitates membrane fusion and delivery of bioactive material to neighboring cells. Small EVs (sEVs; 40-160 nm), also known as exosomes, are generated by two pathways: endosomal sorting complex related transport (ESCRT)-dependent, involving genes such as *TSG101*, and ESCRT-independent that involves *SMPD3*^19,20^. EVs secreted by neoplastic cancer stem and glial cells have been implicated in the reprogramming of non-neoplastic stromal cells in the tumor microenvironment^25^.

Here, we investigated whether *SMPD3* regulates IDH-mutant glioma growth via cell autonomous/autocrine signaling, and EV-mediated, non-cell autonomous/paracrine signaling. Analyzing bulk RNA-seq data from The Cancer Genome Atlas (TCGA), we found that *SMPD3* expression correlates positively with longer survival in patients with IDH-A and IDH-O, consistent with *SMPD3* inhibiting tumor growth. Functionally, *SMPD3* knock-down enhanced the proliferation of IDH-O cells in 2D cultures, in cerebral organoid (CO)/IDH-O co-cultures, and in mouse xenografts with evidence of both cell autonomous and non-cell autonomous effects. The paracrine effects were mediated, at least in part, by IDH-O-derived EVs, which were enriched in ribosomal proteins and induced non-neoplastic astrocyte cell death *in vitro*. Finally, scRNA-seq data mining revealed that non-neoplastic oligodendrocytes within IDH-O and IDH-A tumors acquire a unique transcriptomic signature associated with elevated protein translation. Taken together, these data support the tumor suppressor-like functions of *SMPD3* in IDH-mutant, low-grade gliomas, and suggest that these effects are mediated via cell autonomous and non cell autonomous routes.

## RESULTS

### High *SMPD3* expression correlates with longer survival in IDH-mutant gliomas

Since EVs play a crucial role in remodeling the tumor microenvironment ^26^, we reasoned that EV biogenesis may control IDH-mutant glioma tumor growth. Consequently, we investigated whether the expression of EV biogenesis genes correlated with the survival of patients with IDH-mutant lower grade gliomas. Bulk expression data was extracted from TCGA and the expression levels of genes associated with ESCRT-dependent (*TSG101*) ^27^ and ESCRT-independent (*SMPD3*) EV biogenesis were examined. In Kaplan-Meier survival curves, we compared low versus high gene expression using a log-rank test and found that higher *TSG101* and *SMPD3* expression levels correlated with longer survival times when analyzing all IDH-mutant lower grade gliomas together (Fig. 1a). When controlled for specific tumor type, the survival effect of high *SMPD3* expression remained significant for both IDH-A and IDH-O (Fig. 1b), though the effect was more pronounced for patients with IDH-O (Fig. 1c). Since the survival advantage of high *SMPD3* expression was more marked for IDH-O than for IDH-A, and because *SMPD3* transcripts levels were also higher in IDH-O compared to IDH-A, we chose to focus the remainder of our investigations on IDH-O.

**Figure 1.**
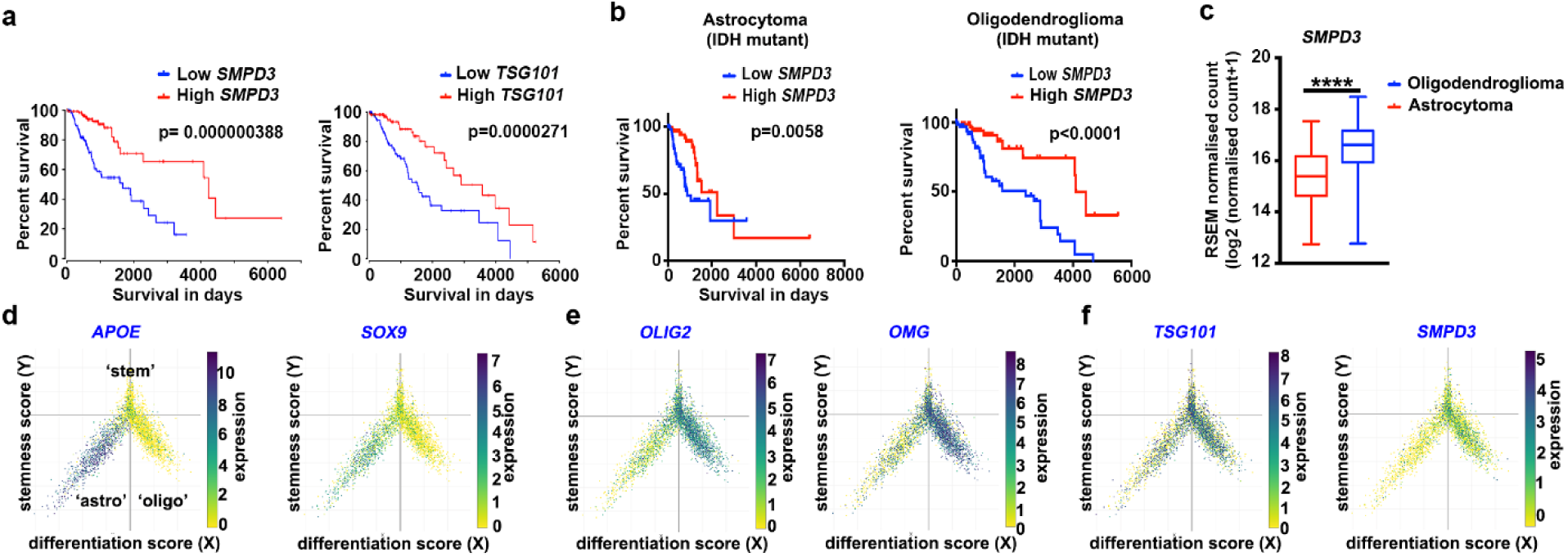
The Cancer Genome Atlas Database (TCGA) analysis showing correlation between *SMPD3* expression and survival of patients with low-grade gliomas. **a-c** Kaplan-Meier survival curves correlating low-grade glioma patient survival with *SMPD3* and *TSG10* expression levels (Log-rank (Mantel-Cox) test; P< 0.0001, **a**); correlating *SMPD3* expression and survival of patients with astrocytoma (Log-rank (Mantel-Cox) test; χ2=7.601; *p=0.0058*; high *SMPD3*>15.81 (n=66); low *SMPD3*<15.0 (n=65), **b**), and correlating *SMPD3* expression and survival of patients with IDH-O (Log-rank (Mantel-Cox) test; χ2=15.27; *p<0.0001*; high *SMPD3*≥16.96 (n=66); low *SMPD3*≤16.28 (n=66), **b**). *SMPD3* levels are higher in oligodendroglioma (IDH-O) patients compared to astrocytoma (IDH-A) (\*\*\*\**p<0.0001*, **c**). **d-f** single cell (sc) RNA-seq showing the expression of genes associated with the astrocytic lineage (*APOE, SOX9*, **d**); oligodendrocytic lineage (*OLIG2, OMG*, **e**); and extracellular vesicle biogenesis gene (*SMPD3, TSG101*, **f**) in malignant IDH-O tumour cells. In each graph, the x axis represents the differentiation score (*i.e.,* glial lineage) and y axis is the stemness scores.

The TCGA data we analyzed was generated from bulk RNA-seq of tumors, such that the gene expression signatures average transcript counts in neoplastic and non-neoplastic cells. To determine whether *SMPD3* was specifically expressed in neoplastic cells in IDH-O, we queried a single cell (sc) RNA-seq dataset in which 1p/19q co-deletion, IDH1/2 mutations, and copy number variations (CNVs) were used to distinguish neoplastic cells from non-neoplastic cells in the microenvironment ^28^. Transcriptional signatures of the neoplastic cells clustered into three main cell types; stem cells, astrocytes, and oligodendrocytes, which when plotted based on their ‘stemness’ and ‘differentiation’ scores in developmental trajectory plots, demonstrates the bipotent nature of IDH-O stem cells ^28^ (Fig. 1d,e). We tused this data to assess exosome biogenesis gene expression in IDH-O neoplastic cells, revealing that while *TSG101* is expressed across the differentiation spectrum, *SMPD3* is enriched in cells with an oligodendrocytic gene expression signature (Fig. 1f). These data confirm that IDH-O neoplastic cells express *SMPD3*, and suggest that *SMPD3* might bias IDH-O cells towards oligodendrocytic differentiation.

### *SMPD3* knock-down increases IDH-O tumor cell proliferation *in vitro*

Given its designation as a tumor suppressor gene in other cancers ^13-16,29^, we reasoned that *SMPD3* could influence IDH-O tumor growth. To determine whether *SMPD3* has a cell autonomous role in regulating IDH-O cancer cell proliferation and/or survival, we used two patient-derived cell lines; BT054 and BT088 cells, derived from grade II and grade III IDH-O tumors, respectively ^30^. To knock-down *SMPD3*, BT088 and BT054 cells were transduced with lentiviral constructs containing one of four sh*SMPD3* sequences (A-D) or an shScrambled (shScr) control sequence, all of which co-expressed GFP (Fig. 2a). nSMase2 protein expression was detected in both BT054 and BT088 tumour cells, as revealed by co-immunolabeling with nSMase2 and SOX10, a marker of glial tumours ^31^ (Fig. 2b). The ability of sh*SMPD3*-B-D sequences to knock-down endogenous SMPD3 protein expression was confirmed in BT088 cells by western blot (Fig. 2c). As expected, BT088 cells expressing sh*SMPD3* B-D variants also secreted fewer CD9^+^ EV particles compared to shScr control transduced cells (Fig. 2d).

**Figure 2.**
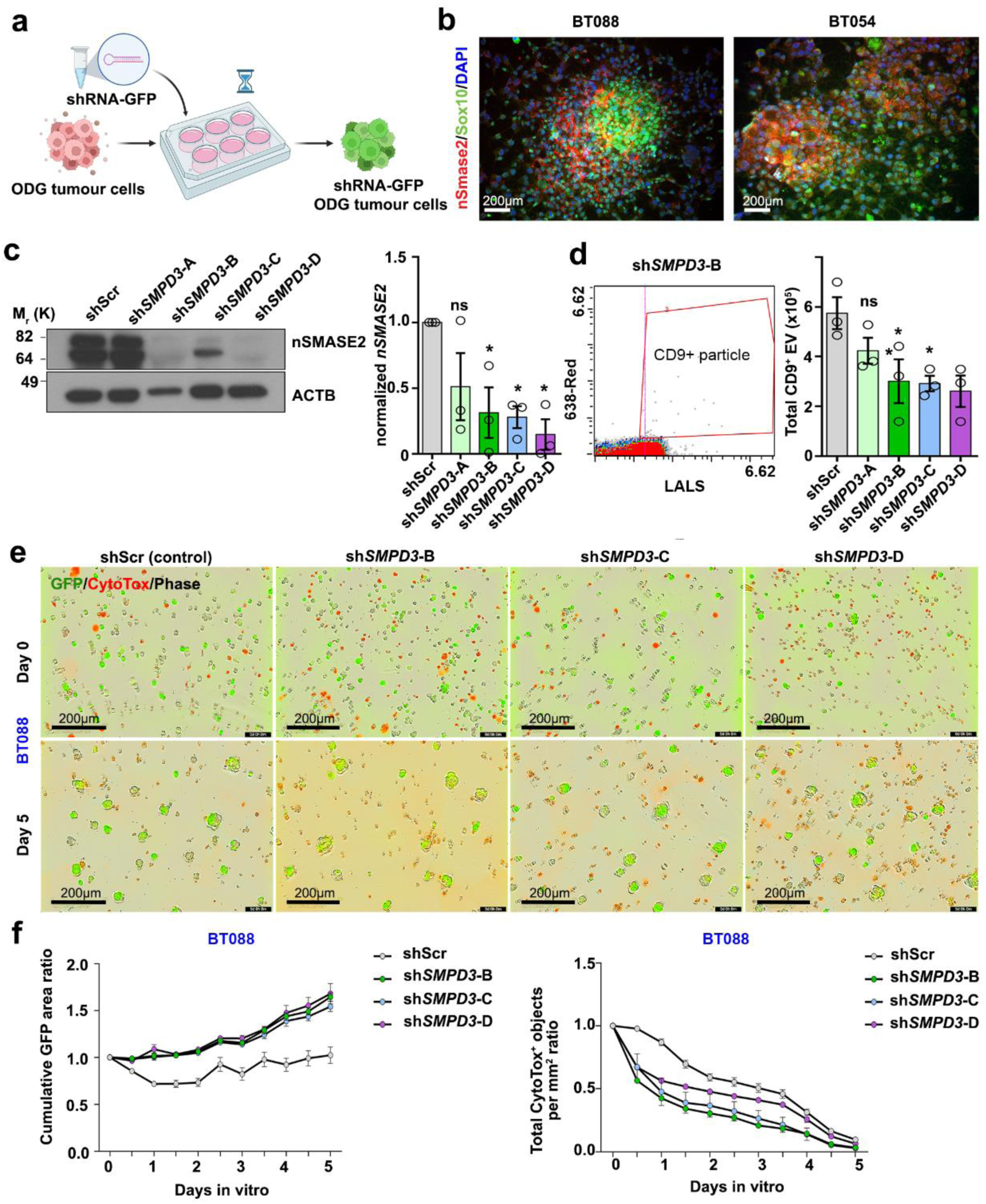
*SMPD3* knock-down promotes IDH-O tumor cell growth *in vitro.* **a** Schematic representing experimental setup to knock down *SMPD3* in IDH-O tumor lines. **b** BT088 and BT054 cells co-immunostained with nSMase2 (red) and SOX10 (Green). Blue is DAPI nuclear counterstain. **c** Western blot showing relative nSMase2 expression levels in shScr and sh*SMPD3* knocked-down (variants A, B, C, D) BT088 cells. **d** Nanoscale flow cytometry showing total CD9^+^ EV particles in shScr and sh*SMPD3* BT088 cells (variants A, B, C, D). **e** Live cell imaging of GFP^+^ shScr and sh*SMPD3* (variants B, C, D)-transduced BT088 cells on day 0 and day 5. Cells were grown with Cytotox red dye. **f** Quantitation of BT088 cell growth by measuring the cumulative area covered by GFP^+^ cells and cell death measured by quantifying the total Cytotox^+^ objects ratio, normalized to day 0. Scale bars: 200 μm.

To determine whether *SMPD3* regulates IDH-O cell proliferation and survival, we used live cell imaging. The cumulative area covered by GFP^+^ cells (normalized to day 0) was measured to monitor cell proliferation and incorporation of Cytotox^+^ dye, a fluorescent dye that enters dying cells due to their increased permeability, was assayed to measure cell death. BT088 cells expressing sh*SMPD3* B-D variants proliferated more rapidly than shScr control cells over 5 days *in vitro* (DIV), with doubling times of approximately 160-192 hrs compared to 800 hrs for shScr control (Fig. 2e,f). Cytotox dye incorporation was lower in the sh*SMPD3* BT088 lines compared to control in the first day of culture, but was not different by the end of the five-day period (Fig. 2e,f). Similar trends were observed after the knock-down of *SMPD3* in BT054 cells, with enhanced proliferation over 5 DIV, without a change in the rate of cell death compared to shScr control lines (Supplementary Fig. 1a,b). Thus, the main effect of *SMPD3* knock-down in IDH-O tumor cells is to enhance cell proliferation, resulting in faster rates of cell expansion *in vitro*.

### *SMPD3* knock-down increases IDH-O invasiveness and proliferation in human cerebral organoids

In patients, brain tumor growth depends on the ability of tumor cells to invade a 3D extracellular matrix (ECM). To provide further support for the role of *SMPD3* as a cell-autonomous regulator of IDH-O proliferation and invasion, we used a 3D co-culture model that better mimics cell-cell and cell-ECM interactions that normally occur in the brain tumor microenvironment ^32^. To monitor the impact of *SMPD3* knock-down, IDH-O tumor cells were co-cultured with COs that were generated from human embryonic stem cells (hESCs) using a modified Lancaster protocol ^33^ (Fig. 3a; Supplementary Fig. 2a). These COs were embedded in Matrigel, an ECM source, but also expressed genes that support their own ECM production ^34^, as evidenced by COL4 and LAM deposition (Supplementary Fig. 2d). Derivative COs were comprised of neural rosettes containing SOX2^+^ neural progenitor cells and MAP2^+^ neurons (Fig. 3b; Supplementary Fig. 2b,c) ^33,34^. While BT088 cells infiltrated into and proliferated within the COs (Fig. 3c-e), BT054 cells instead remained on the surface of the COs, at least with the specific CO protocol and stage used in this study (Supplementary Fig. 2b-d). This finding is consistent with the inability of BT054 cells to thrive in xenotransplants in the murine brain ^30^.

**Figure 3.**
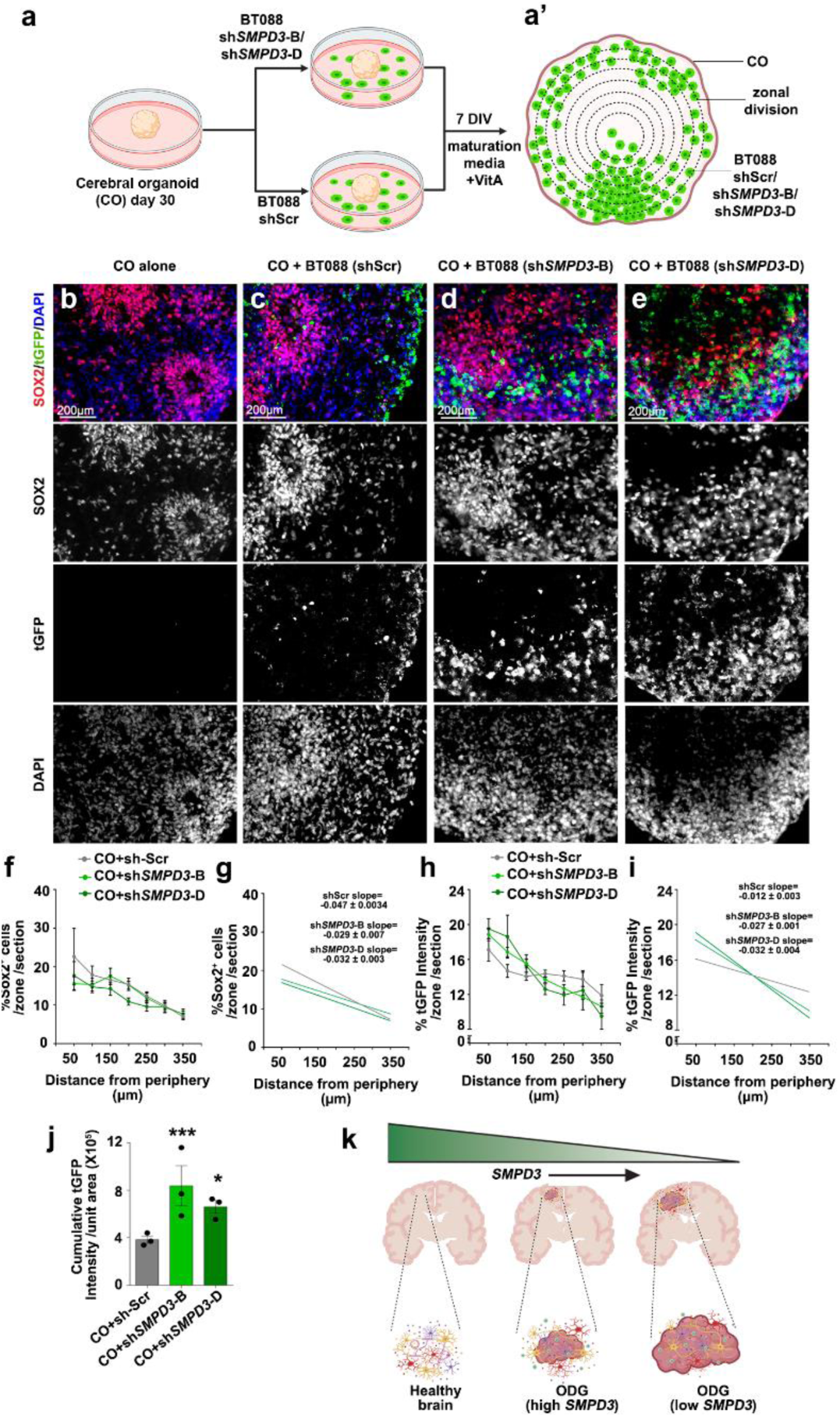
*SMPD3* knock-down facilitates oligodendroglioma cell line growth in human cerebral organoids (COs). **a,a’** Schematic of CO-shScr and sh*SMPD3* BT088 tumor cell co-cultures (**a**), depicting quantitation method and zone division (**a’**). **b-e** COs grown alone (**b**) or in BT088 co-cultures (**c-e**), immunolabeled for SOX2 (red) and turbo GFP (t-GFP; green). Blue is DAPI nuclear counterstain. **f** Percentage of SOX2 ^+^ cells in each of 7 zones. **g** Best-fit lines from (**f**) plotted for the three conditions. shScr: slope= -0.04789 ± 0.003408; sh*SMPD3*-B: slope= -0.02941 ± 0.007366, *p= 0.1234;* sh*SMPD3*-D: slope= -0.03215 ± 0.003388, *p= 0.2205*. **h** Percentage of t-GFP intensity/zone/section. **i** Lines of best-fit (**h**) plotted for shScr: slope= - 0.01259 ± 0.003080; sh*SMPD3*-B: slope= -0.02711 ± 0.001588, *p= 0.003646*; sh*SMPD3*-D: slope= -0.03269 ± 0.004513, *p= 0.000118.* **j** Cumulative t-GFP intensity/unit area in shScr and sh*SMPD3*-COs. **k** Summary of major findings. *\*, p < 0.05; ***, p < 0.001*. Scale bars: 200 µm.

Since BT054 cells did not invade and proliferate in the COs (Supplementary Fig. 2), to evaluate the effects of *SMPD3* knock-down on IDH-O cell proliferation in COs, we used only BT088 cells. 30-day-old COs were cultured either alone or together with BT088 cell lines engineered to express GFP and shScr, sh*SMPD3*-B or sh*SMPD3*-D for 7 days *in vitro* (DIV) (Fig. 3a, a’). We first asked whether the distribution of SOX2^+^ rosettes was perturbed when CO were cultured with tumor cells for 7 DIV (Fig. 3b-e). SOX2^+^ cells were quantified in seven zones from periphery to the core (zone width =50 µm; Fig. 3f). The negative slope of the lines of best-fit suggested that there was a biased distribution of neural rosettes in the CO periphery. However, the slopes were similar for shScr and sh*SMPD3* B-D co-cultures, indicating that IDH-O cells did not alter neural progenitor cell organization (Fig. 3f, g).

Next, we examined the proliferation and distribution of turbo-GFP (tGFP)-labeled IDH-O cells within COs. tGFP^+^ tumor cells were detected in all CO co-cultures (N=3 for each condition; Fig. 3c-e). For sh*SMPD3*-B and sh*SMPD3*-D BT088 lines, more tGFP^+^ cells were detected, aggregating in the CO periphery, where SOX2^+^ neural rosettes preferentially form, compared to shScr controls (Fig. 3c-e), as revealed by the significant difference in slopes of the lines of best-fit (Fig. 3h,i). Moreover, BT088 *SMPD3* knock-down lines had overall more tGFP^+^ cells in the CO compared to shScr (Fig. 3j), consistent with the proliferation advantage of sh*SMPD3* knock-down cells in 2D cultures (Fig. 2). Thus, *SMPD3* knock-down facilitates IDH-O tumor cell proliferation and invasion in a 3D co-culture model that is more reminiscent of the normal tumor microenvironment (Fig. 3k).

### *SMPD3* knock-down facilitates IDH-O tumor growth *in vivo*

Since *SMPD3* knock-down increases IDH-O tumor cell proliferation in 2D and 3D cultures, the knockdown cells would be expected to have greater tumorigenicity *in vivo*. Due to the inability of BT054 cells to form tumors in xenografts ^30^, we again only used BT088 cells, isolated from a higher-grade III tumor, which retains the ability to form tumors after xenografting in immunocompromised mice^30^. To determine whether *SMPD3* knock-down influenced tumor establishment and survival of animals *in vivo,* we generated *Smpd3* knock-down (sh*Smpd3*) and shScr control BT088 cell lines. *SMPD3* knock-down was validated by western blot and sh*Smpd3* BT088 cells were shown to form more and larger tumorspheres than the shScr control cells, consistent with a proliferation advantage (Supplementary Fig. 3a-d).

BT088 cells transduced with shScr (N=8 animals) or sh*Smpd3* (N=8 animals) were transplanted orthotopically into the striatum of NOD *scid* Gamma (NSG) mice (Fig. 4a). Engrafted cells expressed human nuclear antigen (HNA) and nSMase2 (Fig. 4b). By day 154 post engraftment, all mice xenografted with sh*Smpd3* BT088 cells were symptomatic and reached the humane endpoint, whereas two mice xenografted with shScr BT088 cells were still alive after 6 months (181 days), our experimental endpoint (Fig. 4c). Survival curves showed significant differences between the two groups, with shorter survival for animals carrying sh*Smpd3* BT088 cells compared to those with shScr (Mantel-Cox log rank test *p=0.0074*, Fig. 4c).

**Figure 4.**
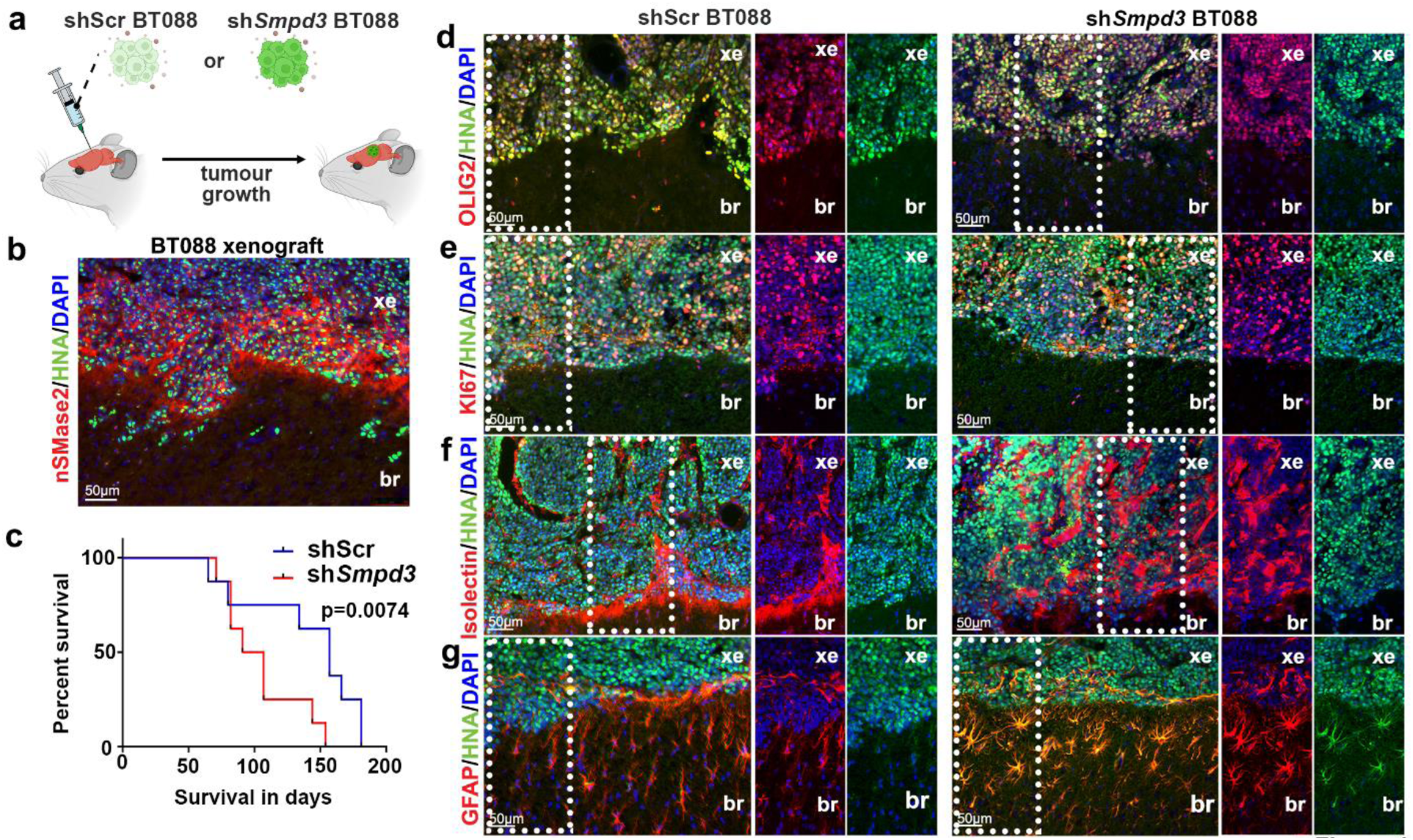
Knock-down of *SMPD3* facilitates IDH-O tumor growth *in vivo*. **a** Experimental protocol followed to xenograft *SMPD3* knock-down BT088 IDH-O tumor cells in cerebral cortices of immunocompromised mice. **b** BT088 tumor xenograft labeled with nSMase2 (red) and human nuclear antigen (HNA, green). Blue is DAPI nuclear counterstain. **c** Kaplan–Meier survival curve associated with shScr and sh*SMPD3* BT088 cell xenografts. Log-rank (Mantel-Cox) test; χ^2^= 7.162; *p=0.0074*. **d**-**g** Immunostaining of shScr (left column) and sh*SMPD3* BT088 xenograft (right column) sections for HNA (green) and OLIG2 (red, **d**), KI67 (red, **e**), Isolectin (red, **f**), and GFAP (red, **g**). Blue is DAPI nuclear counterstain. Insets present split channel images of regions marked by a dotted box. br: normal brain. xe: xenograft. Scale bars are 50μm.

Considering the different survival rates of xenografted mice, we next asked how IDH-O tumor growth impacted other cells in the tumor microenvironments. To characterize the tumor masses associated with *SMPD3* knock-down, we co-immunostained sections with HNA to identify human tumor cells, OLIG2, an IDH-O tumor and oligodendrocyte lineage marker, GFAP, an astrocytic marker, Isolectin, a vascular marker, and KI67, to mark proliferating cells (Fig. 4d-g). Both BT088 shScr and sh*Smpd3* cells co-expressed HNA and OLIG2 (Fig. 4d) and a subset of HNA^+^ tumor cells were KI67^+^ in both tumor types (Fig. 4e). However, the two tumor types differed as sh*Smpd3* tumors appeared to be more vascularized than control shScr tumors, with dense Isolectin staining (Fig. 4f), phenocopying the increase vascular endothelial proliferation that is associated with higher-grade gliomas in patients ^35^. In addition, GFAP^+^ cells bearing morphologic features of reactive astrocytes (hypertrophy of soma and processes, coarser and enlarged processes, more abundant cytoplasm) were more numerous in sh*Smpd3* tumor margins (Fig. 4g), suggestive of glial reactivity ^36^. Together, these results suggest that *SMPD3* knockdown in BT088 cells promotes more aggressive tumor growth in mice, and suggest that non-neoplastic cells in the tumor microenvironment are also impacted.

### IDH-O-secreted EVs induce human astrocyte cell death *in vitro*

The tumor xenograft data suggested that IDH-O tumor cells interact with non-neoplastic cells in the tumor microenvironment. To begin to assess the paracrine effects of IDH-O tumor cells, we focused on EV-mediated communication given the central role that *SMPD3* plays in EV biogenesis ^19,20^. To confirm that BT088 and BT054 cells produce EVs, we first used scanning electron microscopy (SEM), revealing that spherical-shaped, vesicle-like structures with a high polydispersity formed on the cell membranes of both cell lines (Supplementary Fig. 4a). To isolate these vesicles for more direct visualization, we collected conditioned media from each cell line, and used sequential ultracentrifugation as an enrichment method for EVs (Supplementary Fig. 4b) ^27^. Sampling of BT088 cell-derived EVs by transmission electron microscopy (TEM) confirmed their characteristic lipid-bilayer-enclosed morphology (Supplementary Fig. 4c). Nanosight tracking analysis (NTA) showed that both BT054 and BT088 cells produced EVs within the exosome size range (*i.e.,* 40-160 nm ^37^, Supplementary Fig. 4d).

Before testing the bioactivity of BT088 and BT054-derived EVs, we first validated that our preparation method enriched for EVs. Using nanoscale-flow cytometry, 13.7±2.0% of BT088 EVs expressed CD9, a marker of a subset of EVs ^38^ (Supplementary Fig. 4e). Furthermore, western blot analyses of the BT088 cell-derived EV preparations showed the presence of EV-associated proteins (ALIX, CD9^38^, CETP^39^, and FLOT1^40^) and the absence of proteins associated with other organelles, including the endoplasmic reticulum (Calreticulin, Calnexin), mitochondria (VDAC), Golgi bodies (GM130), and peroxisomes (PEX5) (Supplementary Fig. 4f). Since the sequential centrifugation method of EV isolation is sedimentation-based, and thus, can isolate non-vesicular components as well ^41^, we also used density gradient ultracentrifugation for size fractionation to examine EV purity. BT088 EVs were loaded onto a discontinuous Optiprep^TM^ gradient, and after ultracentrifugation, eight fractions were collected and analyzed by western blotting and NTA. EV-associated markers (ALIX, CD9, CETP) were detected in fractions 4 and 5, with a density of 1.08-1.09 g/cm^3^, while markers of other organelles (VDAC, Calnexin) were absent in these fractions (Supplementary Fig. 4g). Of note, CETP was also detected in low-density fractions 1 through 3 (0.95-1.07 g/cm^3^), which contain non-vesicular low-density lipoproteins to which CETP associates ^42^ (Supplementary Fig. 4g). Finally, NTA confirmed that EV particles in layer 4 were predominantly in the exosome size range (Supplementary Fig. 4h). Thus, the EV pellets collected by ultracentrifugation predominantly includes exosomes.

Once the identity of IDH-O cells-derived EVs was confirmed, we investigated the bioactivity of cargo transferred by these EVs by studying their effects on normal human astrocyte proliferation and survival *in vitro*. To do so, normal human astrocytes were plated either in fresh media (FM) or in FM to which isolated EVs from BT054 and BT088 cell were added (FM+EV) (Fig. 5a,b). Using real-time live-cell imaging, cell proliferation and death were monitored over 5 days (Fig. 5c,d). Based on total phase area, FM supported the exponential proliferation of astrocytes, whereas FM supplemented with EVs derived from either cell line reduced (BT054 cells) or prevented (BT088 cells) astrocyte expansion (Fig. 5e,f). To assess cell death, we incorporated the fluorescent Cytotox dye and measured total Cytotox^+^ objects per mm^2^ ratio. Astrocytes grown in FM+EVs from BT054 and BT088 cells incorporated Cytotox dye over the 5 days in culture, with cell death more rapid in the presence of BT088-derived exosomes (Fig. 5e,f). Thus, our in vitro data suggest that BT054 and BT088 cells secrete bioactive exosomes that negatively affect astrocyte proliferation and survival.

**Figure 5.**
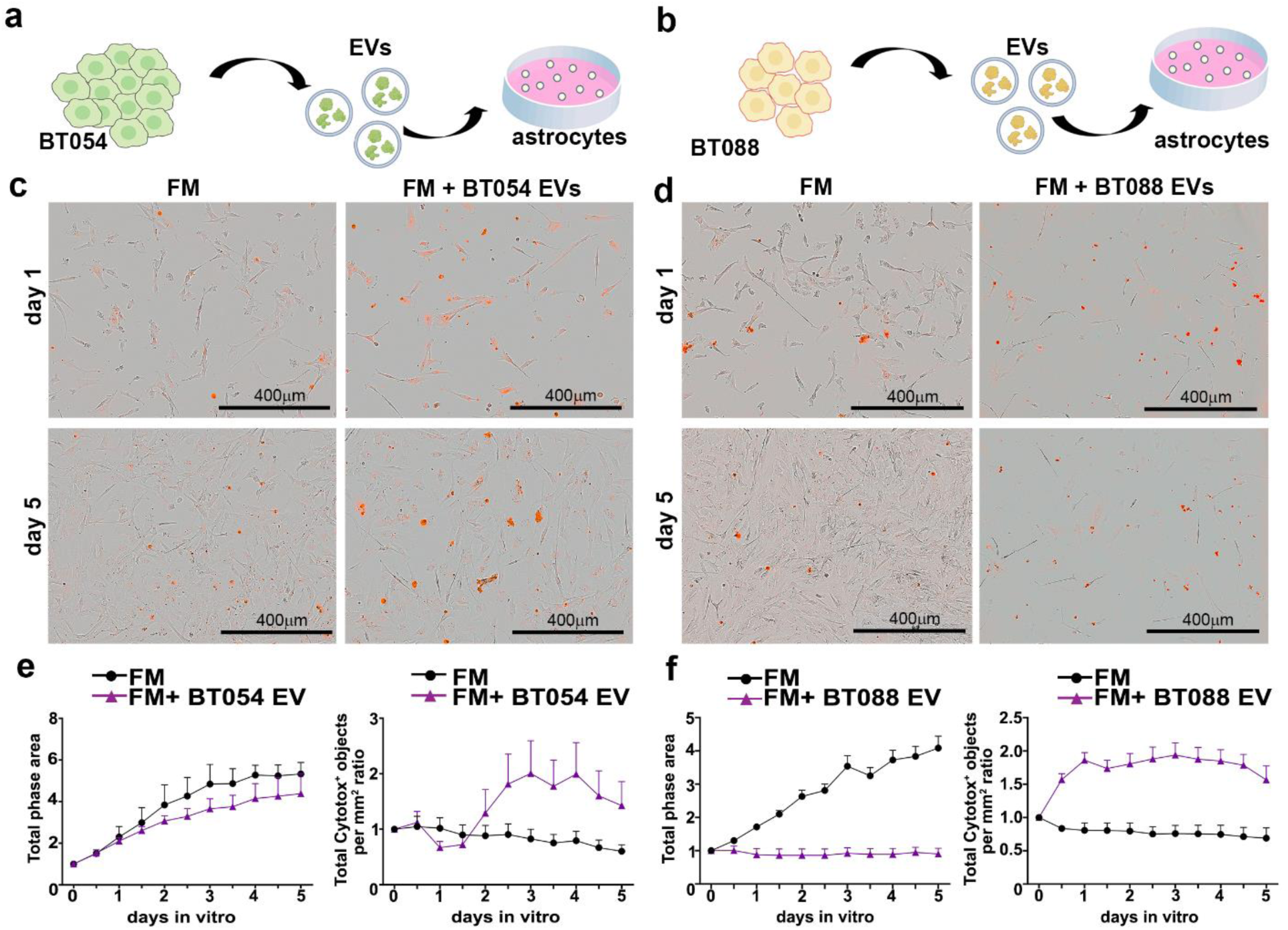
IDH-O-derived extracellular vesicles limit astrocyte growth *in vitro*. **a, b** Schematic representing the experimental setup followed to assess the effect of IDH-O derived EVs on the normal human astrocytes. **c,d** Human astrocytes grown in fresh media (FM) or FM supplemented with BT054 EVs (**c**) or BT088 EVs (**c**), imaged on day 1 and day 5 of growth. Media was supplemented with Cytotox red dye to monitor cell death. **e, f** Quantification of growth and death rates of human astrocytes cultured in FM or FM+BT054 EVs (**e**) or in FM or FM+BT088 EVs (**f**). Cell growth was monitored by measuring the total phase area normalized to day 0. Cell death was monitored by measuring the area covered by Cytotox^+^ cells normalized to day 0. Bars represent means ± s.e.m..

### Evidence for cellular cross-talk within the tumor microenvironment in IDH-O tumors

The tumor xenograft data suggested that IDH-O tumor cells interact with non-neoplastic cells in the tumor microenvironment (*e.g.* increased vascularization and the presence of more reactive astrocytes in the tumor margins). We therefore asked whether we could detect changes in non-neoplastic cells in the tumor microenvironment from surgical resection specimens from patients with grade II and grade III IDH-O and IDH-A tumors (Supplementary Table 1a,b). The identity of neoplastic tumor cells in paraffin-embedded tumor resections was confirmed by IDH1-R132H (hereafter IDHm) immunostaining ^43^ and co-labeling with OLIG2 ^44^, a marker of IDH-O tumor cells and non-neoplastic oligodendrocytes (Supplementary Fig. 5a,b). As a first assessment of tumor microenvironment interactions, we examined proliferation and apoptosis rates, rare events in healthy adult brains ^45,46^. Based on KI67 expression, proliferation rates were lower in both IDH-O and IDH-A tumors compared to five IDH1-mutant GBM and five IDH1-wildtype GBMs (Supplementary Fig. 6a-d). In addition, of the KI67^+^ proliferating cells detected in IDH-O tumors, a surprisingly large fraction was IDHm^-^, and hence, categorized as non-neoplastic cells (65.97%; p<0.0001; Fig. 6a,b). In contrast, the overall percentage of CC3^+^ apoptotic cells did not differ across glioma subtypes (Supplementary Fig. 6e-h). Moreover, similar numbers of IDH-mutant tumor cells (43.66%) and IDH-wild-type non-neoplastic (56.34%) cells expressed CC3 within IDH-O tumors (Fig. 6c,d; p= 0.3484).

**Figure 6.**
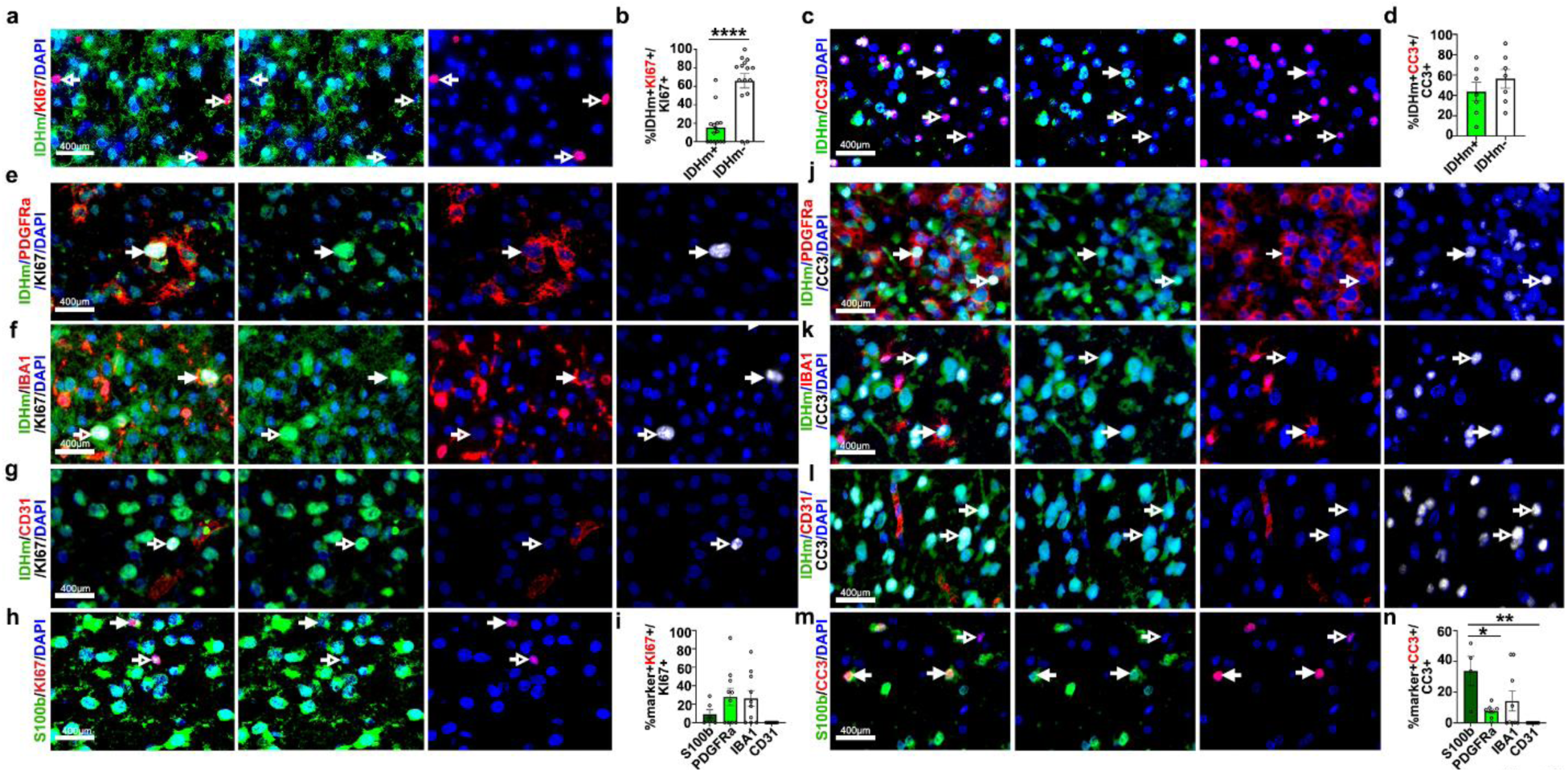
Increased proliferation and cell death (apoptosis) of neoplastic and non-neoplastic cells in IDH-O microenvironment. **a-d** IDH-O tumor resections immunolabeled with IDHm (green), KI67 (red, **a**) or CC3 (red, **c**). Blue is DAPI nuclear counterstain. **b** Percent proliferative (KI67^+^) IDHm^+^or IDHm^-^ cells normalized to the total number of proliferative cells. **d** Percent apoptotic (CC3^+^) IDHm^+^ or IDHm^-^ cells normalized to the total number of apoptotic cells. **e-g** IDH-O tumor resections co-immunolabeled with IDHm (green), KI67 (white) and cell lineage markers, including oligodendrocyte precursor cells PDGFRα (red, **e**), microglia/macrophage, IBA1 (red, **f**), and endothelial cells CD31 (red, **g**). Blue is DAPI nuclear counterstain. **h** IDH-O tumor resection co-immunolabeled with astrocytic marker S100β (green), and KI67 (red). Blue is DAPI nuclear counterstain, **i** Proliferation (KI67^+^) rates among different cell lineages normalized to total KI67^+^ cells. **j-l** IDH-O tumor resections co-immunolabeled with IDHm (green), CC3 (white), and cell lineage markers, including oligodendrocyte precursor cells PDGFRα (red, **j**), microglia/macrophage, IBA1 (red, **k**), and endothelial cells CD31 (red, **l**). Blue is DAPI nuclear counterstain. **m** IDH-O tumor resection co-immunolabeled with astrocytic marker S100β (green), and CC3 (red). Blue is DAPI nuclear counterstain, **n** Cell death (CC3^+^) rates among different cell lineages normalized to total CC3^+^ cells. Closed arrows represent cells positive for all sets of markers and open arrows show lack of co-localization of all markers. Bars represent mean ± s.e.m.. \**p<0.05, **p<0.01, ****p<0.0001*. Scale bars: 400 μm.

To identify which non-neoplastic cells were proliferating in IDH-mutant gliomas, we focused on IDH-O, and performed triple immunolabeling with IDHm, KI67 or CC3, and cell lineage markers, including astrocyte/oligodendrocyte (S100b), oligodendrocyte precursor cells (OPCs; PDGFRa), microglia/macrophages (IBA1), and endothelial cells (CD31) (Fig. 6e-n). Within IDH-O tumors, with the exception of CD31^+^ endothelial cells, OPCs (27.84%), microglia/macrophages (26.15%) and S100b^+^ glia (9.2%) all exhibited increased proliferation (Fig. 6e-i). Similarly, a large fraction of S100b^+^ glia (33.73%), OPCs (7.6%), and microglia/macrophages (14.27%) underwent apoptosis, whereas apoptotic CC3^+^ endothelial cells were not detected (Fig. 6j-n). Taken together, these data suggest that IDH-O tumor cells influence non-neoplastic glia and microglia in the tumor microenvironment.

### IDH-O tumors induce a ribosomal rich gene signature in non-neoplastic oligodendrocytes

Of the non-neoplastic cells in the glioma microenvironment, microglia and astrocytes have been implicated in tumor invasion, growth, and immune protection ^47^, whereas oligodendrocytes have been understudied. For instance, GBM-associated astrocytes exhibit a distinctive immunosuppressive phenotype and associated transcriptional changes ^48^. Here, we queried whether non-neoplastic glia in IDH-O similarly have an altered transcriptional signature by mining an IDH-O scRNA-seq dataset ^28^. Cells were stratified on uniform manifold approximation and projection (UMAP) plot for dimension reduction, with malignant cells identified by the presence of CNVs (Fig. 7a) ^28^. Of the non-malignant cells, not carrying CNVs, microglia, macrophage, T-cells and oligodendrocytes were annotated based on cell type-specific marker expression (Fig. 7a) ^28^. Given the evidence for increased proliferation and death of oligodendrocytes in patient specimens (Fig. 6), we focused on IDH-O-associated oligodendrocytes, shown as a *MOG*-expressing cell cluster (Fig. 7b), and compared their transcriptional signature to that of normal oligodendrocytes from a healthy mouse brain ^49^. We detected 402 differentially expressed genes (DEGs) with a log-2-fold-change (log2FC) greater than 1 (*i.e.,* a doubling in the original scaling) and with a padj<0.05, corresponding to a false discovery rate cutoff of 0.05 (Supplementary Table 2). Gene set enrichment analysis (GSEA) of up-regulated DEGs in oligodendrocytes from IDH-mutant gliomas revealed an enrichment of terms associated with protein translation (Fig. 7c,d). Amongst the upregulated DEGs in IDH-O-associated oligodendrocytes were direct mediators of protein translation (e.g., *EIF1, SEPP1*) and multiple ribosomal proteins (e.g., *RPS8*) (Fig. 7e-h). A similar enrichment of ribosomal and translation-related genes was observed when comparing the transcriptome of tumor-associated oligodendrocytes in IDH-A ^50^ to ‘normal’ oligodendrocytes (Supplementary Figure 7; Supplementary Table 3). These data suggest that protein translation rates are elevated in IDH-O- and IDH-A-associated oligodendrocytes and are indicative of inferred paracrine signaling from IDH-mutant malignant cells.

**Figure 7.**
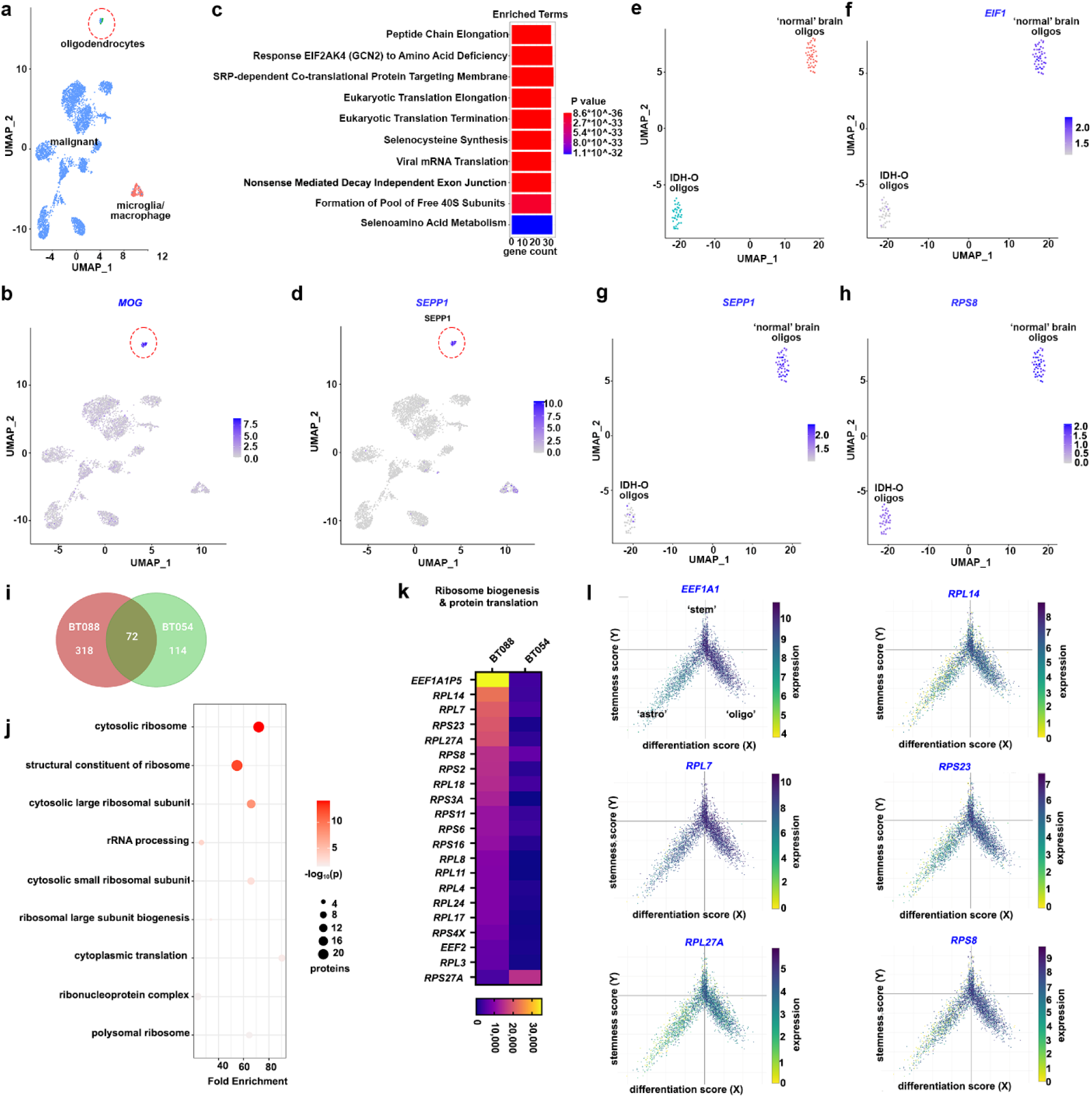
IDH-O-associated oligodendrocytes and IDH-O-secreted EVs are enriched in ribosomal gene/protein signatures. **a** UMAP of scRNA-seq data from IDH-O low grade gliomas ^50^, showing cluster annotation, using CNVs to identify malignant cells, and cell type-specific markers to identify non-neoplastic cells. **b-d** UMAPs on which *MOG* (**b**) and *SEPP1* (**d**) expression were mapped. **c** GSEA of up-regulated genes in IDH-O tumor-associated oligodendrocytes. (**e-h**) Comparisons of gene expression in IDH-O associated oligodendrocytes and ‘normal’ brain oligodendrocytes. **i** Venn diagram of proteins enriched in BT054 and BT088 EVs. **j** Dot plot showing fold enrichment of top 9 identified clusters of proteins involved in ribosome biogenesis and protein translation in KEGG pathway analysis. **k** Heat map of proteins identified in BT054 and BT088 EVs that are associated with ribosome biogenesis and protein translation. **l** Scatter plots of scRNA-seq analysis mapping the expression of top 6 genes expressed in malignant IDH-O tumor cells that are enriched in BT054 and BT088 tumor cell-derived EVs. In each graph, x axis represents the differentiation (lineage) and y axis the stemness scores. Scale bars: 2 μm in **a** (left) and 1 μm in **a** (right); 500 nm in **c** (left); 50 nm in **c** (right).

### IDH-O cells secrete EVs enriched in ribosomal proteins

Finally, we asked how IDH-mutant glioma cells might confer an increased rate of protein translation in non-neoplastic oligodendrocytes. Paracrine effects of cells are mediated through intercellular communication and in part via transfer of EVs ^17,18^. Notably, EVs contain ribosomal proteins that can modify recipient cells, as observed in GBM^51^. Additionally, in this study we found that *SMPD3* knock-down reduces EV biogenesis (Fig. 2c,d). We therefore asked whether the EV proteome might reveal insights into how IDH-O glioma cells impact non-neoplastic cells in the tumor microenvironment. To profile the proteomic fingerprints of EVs secreted by IDH-O malignant cells, we tuned back to BT088 and BT054 cells used throughout this study. EVs were isolated as described (Supplementary Fig. 4), and analyzed using liquid chromatography with tandem mass spectrometry (LC-MS/MS). Of the 390 proteins detected in BT088 EVs and 186 proteins in BT054 EVs, only 72 proteins were common between the two vesiculomes (minimum 2 out of 3 individual replicates; Fig. 7i; Supplementary Table 4). Among the shared proteins were several known EV-associated proteins (*e.g.,* ALIX, CD63), validating the vesicular nature of the EV preparations (Supplementary Table 4). Kyoto Encyclopedia of Gene and Genome (KEGG) pathway analyses of BT088 and BT054 shared vesiculomes showed an enrichment of terms associated with “*cytosolic ribosomes” and “cytoplasmic translation”* (Fig. 7j).

Accordingly, genes encoding for the translation-related and ribosomal proteins were enriched in BT088 and BT054-derived EVs (Fig. 7k). Finally, to confirm that this transcriptomic signature was a feature of IDH-O malignant cells, we mapped the top six-enriched genes from the EV proteome onto the scRNA-seq differentiation trajectory. This analysis revealed an enrichment of transcripts for ribosomal and protein translation-related genes in malignant cells with an oligodendrocytic identity (Fig. 7l). Thus, IDH-O glioma cells secrete EVs that carry ribosomal and translation-related proteins, possibly delivering these proteins to neighboring oligodendrocytes to modulate their cellular state.

## DISCUSSION

IDH-mutant tumors are slow-growing, but the molecular mechanisms underlying their indolent growth remain poorly understood. In this study, our analysis of TCGA revealed a correlation between high *SMPD3* expression with longer survival of patients with IDH-mutant gliomas. Using 2D and 3D cell culture models and xenografts in immunocompromised mice, we found that *SMPD3* functions in an autocrine fashion to limit IDH-O tumor cell proliferation. These results are likely related to the reduced ceramide production caused by *SMPD3* knockdown, and the consequent increase in S1P levels, which induces cell proliferation. In addition, we found a paracrine role for *SMPD3* in IDH-mutant gliomas, likely due to the role of ceramide in driving EV biogenesis ^19,20^. Indeed, EVs derived from IDH-O tumor cell lines induce astrocyte cell death *in vitro*, mimicking the susceptibility of glial cells to undergo apoptosis in biopsies of patient with IDH-O tumors. Furthermore, non-neoplastic oligodendrocytes in IDH-O tumors acquire a transcriptional signature associated with altered ribosomal biogenesis, similar to the enrichment of ribosomal proteins in IDH-O-derived EVs. Taken together, these data support a dual role for *SMPD3* in regulating IDH-O tumor progression, and implicate the encoded nSMase2 enzyme as a potential therapeutic target.

There is growing support for the importance of the sphingolipid rheostat, which balances the lipid second messengers, ceramide (pro-death) and S1P (pro-proliferative), in regulating tumor cell growth ^10^. A previous study revealed that S1P levels are elevated in IDH-wild-type GBM whereas ceramide is elevated in IDH-mutant low-grade gliomas, consistent with these two lipids contributing to the differential growth rates of these tumor types ^11^. Intriguingly, mutant IDH plays a direct role in regulating the sphingolipid rheostat, with inhibitors of the IDH mutant enzyme leading to an increase in S1P levels, which is balanced by a reduction in ceramide ^11^. Our demonstration that the knock-down of *SMPD3*, the critical gene involved in ceramide production, increases the proliferative status of IDH-O cells in vitro and in vivo is in keeping with the sphingolipid rheostat switching in a pro-proliferative (i.e., S1P-producing) direction. Given the importance of this lipid conversion in regulating the growth of multiple tumor types, it is not surprising that drugs targeting the sphingolipid rheostat have now reached clinical trials for glioma patients ^12^.

In this study we also found evidence for non-cell-autonomous effects of *SMPD3* knock-down on the tumor microenvironment, especially impacting non-neoplastic astrocytes. To explain this phenomenon, we focused on the potential impact of exosomes as critical messengers for intercellular communication as they carry macromolecular content that is unique to the cell of origin and deliver this cargo to impact biological and pathological processes in recipient cells ^52,53^. Indeed, exosomes facilitate tumorigenesis by impacting tumor cell growth, angiogenesis, immunity, and tumor metastasis ^52,54,55^. While in many instances, exosomes released within the tumor microenvironment are growth-promoting, anti-tumor factors can also be enclosed in these vesicles ^52^. Accordingly, we show herein that IDH-O cells produce cytotoxic EVs that prevent astrocyte expansion *in vitro.* We also found a correlation between high expression levels of *SMPD3* and *TSG101* with improved patient survival in low-grade glioma patients, suggesting that EV biogenesis via different pathways may play an important role in these tumors. However, it is important to note that increasing EV biogenesis is not always favorable, and depends on the tumor type. For instance, the knock-down of *Rab27a/b*, which also blocks EV secretion, inhibits tumor growth in other brain tumor models ^56,57^. In addition, *Rab27a/b* knock-down in astrocytes, which reduces EV secretion, blocks the metastasis of breast cancer cells to the brain ^56^. Finally, the knock-down of *Rab27a/b* in an astrocyte-derived glioma cell line blocked glioma growth *in vivo* in mouse xenografts ^57^. In high-grade GBM, tumor cells secrete EVs that contain many growth factors (*e.g.,* EGFRvIII) that could promote glioma growth and progression ^58,59^. A likely cause of these different effects is that EV cargo differs based on the cell of origin ^60^.

An important question for future study is what are the bioactive molecules in IDH-O-derived EVs. EV loading of macromolecules occurs either passively or by specific routes that enrich for precise, cell type-specific cargo within EVs, with the vesicular proteome providing a molecular fingerprint of cell identity^61^. GBM-EVs contain bioactive macromolecules that promote tumor growth, including proteins (EGFRvIII, CLIC1), mRNAs (*Annexin A2*), microRNAs (*miR-10b*, *miR-21*, *miR-221*), and long non-coding RNAs (lncRNA) (antisense transcript of AHIF) ^59,62-67,68,69,70^. Several of these molecules serve as prognostic tools for clinical stage and therapeutic response (*e.g.,* CLIC1, *miR-21* ^5,71^). GBM EVs also influence cells in the tumor microenvironment by reducing expression of the tumor suppressor gene *TP53* and increasing expression of the *MYC* oncogene ^72^. In contrast, studies on EVs secreted by low-grade gliomas are in their infancy. IDH-O EVs have been isolated from a single patient-derived IDH-O cell line, demonstrating effects on migratory capacity of peripheral blood mononuclear cells ^73^. Other ‘IDH-O’ studies used G26/24 cells, a mouse line classified as IDH-O-like based on histology, ^74^, but these cells do not model human disease genetically and their relevance to human IDH-O remains unclear. Nevertheless, G26/24-EVs have cytotoxic effects on neurons and astrocytes *in vitro* ^75,76^. EVs isolated from G26/24 ‘IDH-O’ cells exert a cytotoxic effect on neurons and astrocytes *in vitro,* acting via Fas ligand (Fas-L) and tumor necrosis factor-related apoptosis-inducing ligand (TRAIL), respectively ^75,76^. We similarly found using patient-derived BT054 and BT088 IDH-O cell lines that derivative EVs have cytotoxic/cytostatic effects *in vitro*, but we did not detect Fas-L and TRAIL in BT054 and BT088 vesiculomes. Instead, the vesiculomes of BT054 and BT088 cells reveal a common proteomic signature that is associated with ribosomal biogenesis and protein translation, especially in the more aggressive BT088 cells.

Intriguingly, ribosome and protein synthesis genes are expressed at elevated levels in metastatic, circulating breast cancer cells^77^, promote glioma cell plasticity and stemness ^51^, and are associated with worse clinical outcomes in gliomas^77^. Ribosomal proteins are also commonly deposited in EVs in tumor cells, and have been shown to be directly involved in some cases in reprogramming cells in the tumor microenvironment^25^. The identified ribosomal proteins in BT054 and BT088 exosomes have been linked to tumor development and progression. For instance, RPL14 has been implicated in esophageal squamous cell carcinoma ^78^ and cervical cancer ^79^, RPL18 in colorectal cancer ^80^, RPL27a in triple-negative breast cancer ^81^ and hepatocellular carcinoma ^82^, and RPS3a in hepatocellular carcinoma ^83^. Moreover, the high expression level of RPS11 was shown to be associated with poor prognosis and patient survival in primary and secondary glioblastoma ^84^. RPS3a and RPL7 together regulate apoptosis in NIH 3T3 ^85^ and Jurkat T lymphoma cell lines ^86^.

While most high-throughput screens have focused on identifying small molecules that block EV secretion, which could be useful for GBM tumors ^87^, there have also been recent screens for small molecules that induce EV secretion ^88^. Interestingly, N-methyl dopamine and norepinephrine activate nSMase2 to increase EV production in mesenchymal stem cells without increasing cell number, a strategy that is being developed to enhance the regenerative potential of MSC-derived EVs ^88^. Testing whether these drugs increase IDH-O EV production, and the associated cytotoxicity, would be of interest in the future. In summary, we have uncovered *SMPD3*-mediated exosome biogenesis as a critical mechanism underlying IDH-O tumor growth and cell interactions with the tumor microenvironment. Controlling the growth of malignant IDH-O cells by manipulating *SMPD3*-regulated pathways may have repercussions that extend beyond survivorship, including the reduction of neurological side effects such as epilepsy and neurocognitive decline that reduce life quality ^89^.

## METHODS

### Animals

8-10-week-old female NOD *scid* Gamma mice (NSG; OD.Cg-Prkdc*^scid^Il2rg^tm1Wjl^*/SzJ; N=8) were purchased from the Animal Research Center (ARC), UHN, Toronto, Canada and housed at the Princess Margaret animal breeding facility, where xenograft experiments were performed, as approved by the UHN Animal Care Committee (protocol 3499.16.2 to VAW).

### Patient-derived formalin-fixed paraffin-embedded (FFPE) tissue resections

Specimen slides of tumor biopsies from patients with IDH mutant IDH-O (WHO Grade II, III) and IDH-mutant anaplastic astrocytoma, IDH-mutant GBM, and IDH-wildtype GBM were obtained from the pathology archives at the Calgary Laboratory Services and Clark Smith Brain Tumor Bank at the University of Calgary (by JAC). Samples were fixed in formalin and embedded in paraffin. The approval to use FFPE tissue specimens was obtained from Calgary Laboratory Services and the Calgary Health Region Ethics Board (the University of Calgary Conjoint Health Research Ethics Board to JAC (HREB #2875 and #24993).

### Cell lines and cultures

BT054 and BT088 lines were collected and approved under protocols from the Health Research Ethics Board of Alberta to JAC (HREBA.CC-16-0762 and HREBA.CC-16-0154). BT054 and BT088 cell line use was approved by the SRI Ethics Board (REB) to CS (PIN:SUN-2614). Cells were maintained in tumor complete media composed of Neurocult NS-A proliferation media (Human; Stem Cell Technologies; cat# 5751) supplemented with human epidermal growth factor (hEGF; 20ng/mL; PeproTech; cat# 100-15), human basic fibroblast growth factor (β-FGF; 20ng/mL; PeproTech; cat# 100-18B), heparin (2ug/mL; Stem Cell Technologies; cat# 07980), and antibiotic-antimycotic solution (0.1%; Wisent; cat# 450-115-EL). BT054 and BT088 cells were cultured as adherent cells on poly-D-lysine: laminin coated tissue culture plates or flasks maintained in a humidified 5% CO^2^ incubator. Cells were dissociated and passaged using Accutase (Stem Cell Tech; cat# 07920) according to the manufacturer’s instructions.

### Conditioned media and EV isolation

1x10^6^ BT054 or BT088 cells were seeded in 11mL media and the conditioned media (CM) was collected after 24 hours. CM was centrifuged at 300 x g for 5 min to remove cells, 2000 x g for 10 min to remove cellular debris and 10,000 x g for 30 min to remove protein aggregates and smaller debris. CM was then sequentially ultra-centrifuged at 100,000 x g for 2 hours at 4°C in a Beckman Coulter Optima L-100 XP ultracentrifuge using an SW41-Ti rotor and polycarbonate centrifugation tubes (Beckman Coulter, #331372). EV pellets were rinsed with phosphate buffered saline (PBS; ThermoFisher Scientific, #14190144) and centrifuged at 100,000 x g, 4ͦC for 1 hour and resuspended in 50 μl PBS prior to use.

### Astrocyte growth assays

Normal human astrocytes (Lonza, #CC-2565) were cultured in astrocyte growth medium (AGM^TM^ BulletKit^TM^; Lonza, #CC-3186), which was changed daily, and cells were maintained at 37°C in a humidified 5% CO^2^ incubator. Cells were dissociated using 0.125% Trypsin (Wisent) and incubated for 2-3 minutes at 37°C. Dissociated cells were seeded at 3000 cells per well in 6-well plates (Corning) in fresh media (FM) or FM+EVs obtained from either BT054 and BT088 cell cultures (as described in cell lines and cultures) for 5 days in vitro (DIV). Cell growth and death were monitored and images were obtained using the Incucyte S3 Live Cell Imaging System (Essen BioScience).

### Incucyte live cell imaging

An Incucyte S3 Live cell imaging system (Essen BioScience) was used to monitor cell growth and death. For unlabeled and GFP^+^ transfected cells or cell lines, growth rates were measured by monitoring the total phase area confluence or area covered by GFP^+^ cells. For cell death assays, prior to cell seeding, media was supplemented with 0.25 uL/mL of Cytotox dye (Red, #4632 or green, #4633) per well per the manufacturer’s instructions. Phase contrast and red or green fluorescent imaging were taken every 12/24 hours for the cell growth studies and every 4/12 hours for the cell death studies at 10x/2x magnifications. A minimum of 9 images per time point per well were taken at each time point. The Incucyte application’s built-in analyzer algorithm was used to quantify cell growth and death rates.

### Cerebral organoid-BT088 co-cultures

The use of hESCs was approved by the SRI Research Ethics Board (REB) to CS (PIN:SUN-1884). Feeder-free H1 hESCs (WiCell) were cultured on Matrigel in TeSR™-E8™ kit for hESC/hiPSC maintenance (StemCell Tech; #05990). hESCs were used to generate cerebral organoids (COs) using media included in the STEMdiff Cerebral Organoid Kit (StemCell Tech; #08570) and STEMdiff Cerebral Organoid Maturation Kit (StemCell Tech; #08571), with some modifications. Briefly, hESCs were plated in 96-well round-bottom ultra-low attachment plates at 9,000 cells/well in embryoid body (EB) seeding medium. Dual SMAD inhibitors (2μM Dorsomorphin; StemCell Tech; #72102, and 2 μM A83-01; StemCell Tech; #72022) were added to the media until day 5. Newly formed EBs were transferred to 24-well plates containing StemCell Tech CO induction medium. On day 9, EBs with optically translucent edges were embedded in Matrigel and deposited into 6-well ultra-low adherent plate with StemCell Tech expansion medium. From day 5 to day 13, media was supplemented with 1 μM CHIR-99021 (StemCell Tech; #72052) and 1 μM SB-431542 (StemCell Tech; #72232) to support formation of well-defined, polarized neuroepithelia-like structures. On day 13, embedded EBs exhibiting expanded neuroepithelia as budding surfaces were transferred to a 12-well spinning bioreactor (Spin Omega^33^) containing maturation medium in a 37°C incubator. For BT088 co-culture, on day 30, COs were individually transferred to a 24-well plate containing Neurocult NS-A proliferation media (Catalog # 05751, StemCell Tech) with freshly added hFGF2 (20 ng/ml), hEGF (20ng/ml), and heparin (2μg/mL). Subsequently, 10,000 GFP^+^ BT088 cells were added to each well. Plates were incubated for 24 hrs without agitation and on the next day tumor-bearing COs were washed with PBS once and maintained in maturation media on an orbital shaker at 37°C for 7 more days. On day 8, COs were fixed in 4% paraformaldehyde for 45 min, transferred into 30% sucrose overnight, snap frozen in Optimal Cutting Temperature (OCT) for cryosectioning.

### The Cancer Genome Atlas (TCGA) analysis

A Kaplan Meier plot was generated from the lowest and highest quartiles of all low-grade glioma patients for *SMPD3*, involved in ESCRT-independent, and *TSG101*, involved in ESCRT-dependent pathways. Low-grade glioma, IDH mutant astrocytoma, and IDH mutant IDH-O datasets were downloaded from UCSC’s Xena Browser (https://xenabrowser.net/) and gene expression levels were correlated with overall patient survival.

### Immunofluorescence staining

FFPE tumor resection specimens were immunostained using the Opal™ 4-Color Manual IHC Kit (NEL820001KT), as per the manufacturer’s protocol. IF staining of tumor xenografts was performed following paraformaldehyde fixation, OCT embedding, and sectioning of the brains. Section slides were washed and permeabilized in phosphate-buffered saline containing 0.1% Triton X-100 (Sigma, #T8787; PBST) followed by blocking with PBST containing 10% normal horse serum (blocking solution; HS, Wisent, #065-150) for 1 hour at room temperature. Next, the sections were incubated with primary antibodies prepared in the blocking solution overnight at 4°C. Sections were washed with PBST and incubated with the compatible species-specific secondary antibodies diluted in PBST for 1 hour at room temperature. Slides were washed with PBST and counterstained with diluted 4′,6-diamidino-2-phenylindole (DAPI; Invitrogen, #D1306) in PBST at room temperature. Finally, the slides were washed with PBST and mounted with coverslips using AquaPolymount (Polysciences Inc., #18606-20). Primary antibodies used included pERK1/2-T202/Y204 (Cell Signaling, #4370), IDH1 R132H (Dianova, #DIA-H09), PDGFRα (Cell Signaling, #3174), S100b (Sigma, S2532), IBA1 (FUJIFILM Wako, #019-19741), CD31 (Abcam, #), KI67 (Abcam, #ab16667), cleaved caspase 3 (CC3; Abcam, #ab2302), GFAP (Millipore, #MAB360), OLIG2 (Abcam, #ab109816), SOX2 (Abcam, #ab97959), HNA (Millipore, #MAB1281), nSMase2 (Abcam, #ab85017), Isolectin (Sigma, #L2140), turboGFP (OriGene, #TA150041), SOX10 (Santa Cruz, #SC-365692), zsGreen (Takara, #632474), MAP2 (Abcam, #5392), COLL4 (Abcam, #Ab6586), and LAM (Sigma, #L9393). The secondary antibodies used were Alexa 568 donkey anti-rabbit, Alexa 488 donkey anti-rabbit, and Alexa 488 donkey anti-mouse (all from Invitrogen).

### Lentiviral transduction

To knock-down *SMPD3,* BT088 or BT054 cells were transduced with four *SMPD3* human shRNA lentiviral particles (A,B,C,D) (pGFP-c-shLenti-sh*SMPD3*A-D; TL301492VA-D; OriGene) or lentiviral shRNA-Scr control particles (pGFP-C-shLenti-shScrambled; TR30021V; OriGene). Titres ranged from 2.0x10^7 to 3x10^7 TU/ml. Lentiviral particles were added to BT088 or BT054 cell cultures seeded in a poly-D-lysine:laminin pre-coated plate. To induce the expression of *SMPD3*-Luc2 and Luc2 (control), transduced GFP^+^ cells were treated with 2μg/mL Doxycycline Hyclate (Millipore Sigma, #D9891).

### Western blotting

Cells or EV pellets were lysed in the lysis buffer containing a cocktail of protease (1X protease inhibitor complete, 1mM phenylmethylsulfonyl fluoride) and phosphatase (50mM sodium fluoride, 1mM sodium orthovanadate) inhibitors. 10μg of lysate was loaded and run on 10% SDS-PAGE gels for western blot analysis. Transfers to PVDF membranes (#1620177, Biorad) were made in transfer buffer (25 mM Tris, 192 mM glycine, 20% methanol, pH 8.3) at 40V overnight at 4℃. Antibody blocking was in TBST (10 mM Tris, 100 mM NaCl, pH 7.4, 0.1% Tween-20) with 5% (W/V) powdered milk for 1 hour at room temperature, which was also the same buffer used for primary antibodies, which were incubated overnight at 4℃. Primary antibodies used at 1/1000 included Flotilin1 (cell Signalling, #3253), CD9 (Santa Cruz, #sc9148), nSMASE2 (Abcam, #ab85017), CETP (Abcam, #ab2726), ALIX (Cell Signalling, #2171S), GM130 (BD Biosciences, #610822), Calnexin (Abcam, #ab22595), Calreticulin (Abcam, #ab2907), PEX5 (Novus Biologicals, #NBP1-87185), VDAC (Cell Signalling, #4661), and β−actin (Abcam, #ab8227). Membranes were washed in TBST, incubated with 1/10,000 dilutions of horseradish peroxidase (HRP)-coupled secondary antibodies (Anti Rabbit IgG #7074S, Cell Signalling Technology, Anti Mouse IgG #Pierce 31430, ThermoFisher), washed in TBST, and then developed with ECL Plus Western Blotting Reagent (#29018904, GE Healthcare) according to the manufacturer’s instructions. Blots were developed using X-ray film (#1141J52, LabForce). Densitometries were calculated using ImageJ and the average values of normalized expression levels were plotted.

### Mouse tumor xenografts

For the mouse xenograft experiments, PB-CMV-GreenPuro-H1-MCS PiggyBac shRNA Cloning and Expression Vector (SBI System Biosciences, # PBSI505A-1) was used. The vector backbone was linearized with BamH1 and EcoR1 and the following annealed oligonucleotides were cloned into the site: sh*Smpd3*:5’pGATCCCCCTCATCTTCCCATGTTACTTCAAGAGAGTAACATGGGAAGAT GAGGGACGCGTG3’(sense) and 5’pAATTCACGCGTCCCTCATCTTCCCATGTTACTCTC TTGAAGTAACATGGGAAGATGAGGGG 3’ (antisense); and for shScr:5’pGATCCATTCAC TTATCCGCCTCTCCTTCAAGAGAGGAGAGGCGGATAAGTGAATCTCGAGG3’(sense), and5’pGAATTCCTCGAGATTCACTTATCCGCCTCTCCTCTCTTGAAGGAGAGGCGGAT AAGTGAATG -3’ (antisense). The sh*Smpd3* construct targeted the mouse *Smpd3* sequence, and the mouse shRNA target sequence has a 3 bp mismatch upon alignment with the human *Smpd3* sequence. Knock-down was confirmed by western blot (Supplementary Fig. 3). In preparation for xenografting, BT088 cells were nucleofected with each sh*Smpd3* or shScr vectors mixed with Super PiggyBac Transposase expression vector (SBI, # PB210PA-1) in 1:3 PiggyBac: Transposase ratio. To perform nucleofection, 2x10^6^ dissociated cells were suspended in 20μL of P3 reagent with the DNA mix (final concentration of 12μg). Nucleofection was performed using 4D -Nucleofector^™^ X Kit S (Lonza, # V4XP-3032) in nucleofector strips using the CZ167 program. 8-10-week-old female NOD *scid* Gamma mice (NSG; NOD.Cg-Prkdc*^scid^* Il2rg*^tm1Wjl^*/SzJ; Princess Margaret animal breeding facility) were used for tumor xenograft studies. Sh*Smpd3*-GFP and shScr-GFP BT088 cells were injected into the right cerebral hemispheres of 8 mice. The coordinates for implantation were AP-1.0, ML 2.0, and DV 3.0. The xenografted cells developed into tumors, for which the mice were monitored daily. Upon development of terminal symptoms, mice were sacrificed at relative endpoints. Two control mice did not show any terminal symptoms post-engraftment and were humanely sacrificed after 180 days.

### Nanosight tracking analysis (NTA) and nanoscale flow cytometry

NTA was performed using the Malvern NanoSight NS300 at the Structural & Biophysical Core Facility, University of Toronto. EV pellets were collected by sequential centrifugation, resuspended in 200 μl PBS, diluted 1:50 and passed through the Nanosight chamber. NTA data acquisition settings were as follows: camera level 13, acquisition time 3 × 30 seconds with detection threshold 12. Data were analyzed with the NTA3.2 Dev Build 3.2.16 software. For nanoscale flow cytometry, EVs (1 μl in 18 μl sterile water) were incubated with CD9 antibody (1 μl; Santa Cruz, #sc9148) for 30 min at room temperature. Post incubation, EVs were stained with Alexa Fluor 647 Far red secondary antibody (1 μl of 1:20 antibody solution; Invitrogen) for 20 min. Stained EVs were diluted in 500 μl sterile water and quantified on the Nanoscale Flow Cytometer (Apogee Flow Systems Inc). A representative scatterplot of BT088 EVs was plotted for 638-Red (detecting Alexa 647 bound CD9^+^ particles) and long angle light scatters (LALS; for size distribution). EVs were defined as size events greater than 100 nm.

### Scanning electron microscopy (SEM)

BT088 and BT054 cells were cultured on 13 mm coverslips (EM Biosciences) precoated with Poly-O-Lysine–Laminin in a 24-well plate. The samples were then fixed (2% glutaraldehyde in 0.1M sodium cacodylate buffer, pH 7.3; for over 2 hours), rinsed, and dehydrated. Samples were mounted on stubs, gold sputter-coated, and imaged with FEI/PhilipsXL30 scanning electron microscope at 15 kV. Imaging was performed in the Nanoscale Biomedical Imaging Facility, SickKids Research Institute.

### Transmission electron microscopy (TEM)

Before grids were prepared, carbon coated Cu400 TEM grids were glow discharged for 30 seconds (Pelco EasiGlow, Ted Pella Inc.). Then 4 μL of BT088 exosome solution was applied to the grid for 60 seconds before wicking away the excess solution. The grid was washed three times with 4 μL of distilled water and stained with 4 μL of 2% uranyl acetate solution for 30 seconds. Excess uranyl acetate solution was wicked away. The grids were air-dried. Imaging was performed on a Thermo Fisher Scientific Talos L120C TEM operated at 120 kV using a LaB6 filament. Imaging was performed in the Microscopy Imaging Laboratory, at the University of Toronto.

### Density gradient ultracentrifugation

OptiPrep^TM^ (Iodixanol 60% stock solution; Stem Cell Tech, #07820) was diluted with a homogenization solution (0.25M sucrose in 10mM Tris HCl pH7.5) to generate a discontinuous density gradient of 40% (2.5mL), 20% (2.5mL), 10% (2.5 mL), and5% (2 mL). The solutions were carefully pipetted in an ultracentrifuge tube and left undisturbed for over 1 hour. EVs (filtered through a 0.2 μm filter and resuspended in 500 μL PBS) were isolated from BT088 conditioned media by sequential centrifugation, and after the first 100,000 x g spin were loaded onto the gradient. Samples were centrifuged at 100,000 x g for 18 hours. Post centrifugation, 1 mL fractions were pipetted out carefully from the top. Fractions were mixed with PBS and centrifuged at 100,000 x g for 4 hours. EV pellets were resuspended in 50 μL PBS.

### Mass spectrometry

Mass spectrometry was performed on BT088 and BT054 EVs isolated by sequential ultracentrifugation at the SPARC Biocentre-Mass Spectrometry facility at SickKids Research Institute. Mass spectrometry analysis information is provided in Table S2 and data analysis in Table S3. Scaffold data analysis was performed by applying NCBI annotations to all proteins, removing proteins that matched the search term “keratin” and which did not have “Homo sapiens” under the taxonomy heading. Protein filtering thresholds were set at 99.0%, with a minimum number of 2 peptides and a peptide threshold of 95%. The Cytoscape program and ClueGO plugin were used to analyze the EV samples. Gene Ontology (GO) enrichment analysis was performed using *rrvgo* (https://ssayols.github.io/rrvgo/) on the significantly differentially expressed proteins and genes to identify relevant biological, molecular, and cellular processes. And pathway enrichment analysis was performed using the *pathfindR* Bioconductor package (https://github.com/egeulgen/pathfindR).

### scRNA-seq data acquisition and analysis of IDH-O cells

scRNA-seq data profile of 4,347 single cells from six IDH1/2 mutant human oligodendroglioma (i.e., IDH-O) containing 4,044 malignant cells was obtained from the Gene Expression Omnibus (GEO) website under the accession number GSE70630^28^, as well as IDH-mutant glioma under the accession number GSE89567. The scatter plots from scRNA-seq data obtained mapping the expression of queried genes in malignant IDH-O cells were downloaded from the Broad Institute Single Cell Portal via https://singlecell.broadinstitute.org/single_cell/study/SCP12/oligodendroglioma-intra-tumor-heterogeneity.

### scRNA-seq data analysis of oligodendrocytes

The scRNA-seq data of oligodendrocytes from IDH-mutant gliomas was obtained from ^90^ and the Smart-seq2 version of the scRNA-seq data of normal oligodendrocytes from ^49^. The processed data from IDH-mutant gliomas, along with UMAP and tSNE coordinates and cell identities and malignancy based on copy number variants (CNVs) was downloaded at the Broad single cell portal: https://singlecell.broadinstitute.org/single_cell/study/SCP50/single-cell-rna-seq-analysis-of-astrocytoma. https://singlecell.broadinstitute.org/single_cell/study/SCP12/oligodendroglioma-intra-tumor-heterogeneity

For the normal oligodendrocytes, the Single Cell Portal Accession number was SCP425 downloaded from Single Cell Portal (https://portals.broadinstitute.org/single_cell). From all three datasets, oligodendrocyte cells were extracted and processed together using the Seurat v.4.0.1 R package ^91^ using the SCTransform function while regressing out the variance due to mitochondrial RNAs. Differential gene expression analysis was conducted between the oligodendrocyte types using the FindMarkers function with min.pct being 0.1 and the adjusted p-value cutoff of 0.05. GSEA was carried out on the differentially up-regulated genes in the oligodendrocytes from IDH-mutant gliomas with the enrichR R package ^92^ using the “Reactome_2022” database.

### Imaging and figure generation

Images were captured with a Leica DMIL LED or DMRXA2 optical microscope or a Leica DMi8 Inverted Microscope (Leica Microsystems CMS, 11889113) using LasX software. ImageJ software was used for image analysis. For patient tumor analysis, single channel TIFF images from all sample sets were transformed to binary format with mean intensity as the selecting parameter. A fixed minimum and maximum threshold value were determined for each set of images to ensure correct thresholding. Single channel images were merged to generate the merged images, and were analyzed by adjusting the size filter option to count cells manually in each sample set. For cerebral organoid-tumor co-culture assessment, zone-wise cumulative GFP intensity and total number of Sox2^+^ cells were analyzed using ImageJ. The freeline tool was used to mark the periphery of the organoid. Seven zones (width=150 pixels or 50 μm) were constructed mapping the shape of each organoid, spanning from organoid periphery (zone1) to the core(zone7). Cumulative GFP intensity and Sox2^+^ cell counts were assessed per zone. Figures were created using Adobe Photoshop and schematics were generated using BioRender.com.

### Statistics and Reproducibility

Statistical analysis and graphs were generated using GraphPad Prism 6 software. One-way ANOVA with TUKEY post-corrections was used when comparing groups of more than two, while student’s t-test was used when comparing two groups. In primary tumorsphere assays in Supplementary Fig. 3, tumorsphere size was measured using the ruler measurement tool in ImageJ. A minimum of three biological replicates were carried out for all assays. All data are expressed as mean value ± standard error of the mean (s.e.m.). In all experiments, a p-value <0.05 was taken as statistically significant, *\*p < 0.05*, *\*\*p < 0.01*, *\*\*\*p < 0.001*, *\*\*\*\*p<0.0001*.

## ACKNOWLEDGEMENTS

We thank Dr. Gregory Cairncross and Dr. Sam Weiss (University of Calgary) for initially providing the BT088 and BT054 tumor cell lines, Ali Darbandi (TEM), Lindsey Fiddes (SEM), and Greg Wasney (NTA) at the University of Toronto for their technical help, and Itay Tirosh (Weizmann Institute) for assistance accessing scRNA-seq datasets. This project was supported by the International Development Research Centre (IDRC 108875) to CS, Canada First Research Excellence Fund (CFREF) Medicine by Design Cycle 2 to CS and CM, a Cancer Research Society grant to CS and JAC, and a Terry Fox Research Institute New Frontiers Program Project Grant to HSL, IPM and CS. CS holds the Dixon Family Chair in Ophthalmology Research.

## AUTHOR CONTRIBUTIONS

AB, FS, LA: conceptualization, data curation, formal analysis, investigation, methodology, visualization, validation, writing – original draft, writing – review, and editing VC, AES, TO, MJC, STA: data curation, formal analysis, software OP, LV, YT, RI, SS, DZ, LCC, BK: data curation, formal analysis, software TF: methodology, investigation HSL: resources, supervision, writing – review, and editing CMM, IPM, HSL, MB, VAW: resources, supervision, writing – review, and editing JAC, CS: funding acquisition, conceptualization, project administration, resources, supervision, validation, writing – original draft; writing – review and editing

