## Supplemental Data for "*SMPD3* suppresses IDH mutant tumor growth via dual autocrine-paracrine roles"

### SUPPLEMENTARY TABLES

**Supplementary Table 1. Summary of patient tumors analyzed.**

**Supplementary Table 2. DEGs upregulated in IDH-O-associated non-malignant oligodendrocytes vs normal brain oligodendrocytes.**

**Supplementary Table 3. DEGs upregulated in IDH-A-associated non-malignant oligodendrocytes vs normal brain oligodendrocytes.**

**Supplementary Table 4. LC-MS/MS analyses of BT054 and BT088-derived extracellular vesicles.**

### SUPPLEMENTAL FIGURES

**Supplementary Figure 1. *SMPD3* knock-down promotes IDH-O tumor cell growth *in vitro*.**
**a, b** Quantitation of shScr, and sh*SMPD3*-BT054 cells (variants b-d) growth by measuring the cumulative area covered by GFP$^+$ cells (**a**) and cell death measured by quantifying the total Cytotox$^+$ objects ratio (**b**), normalized to day 0.

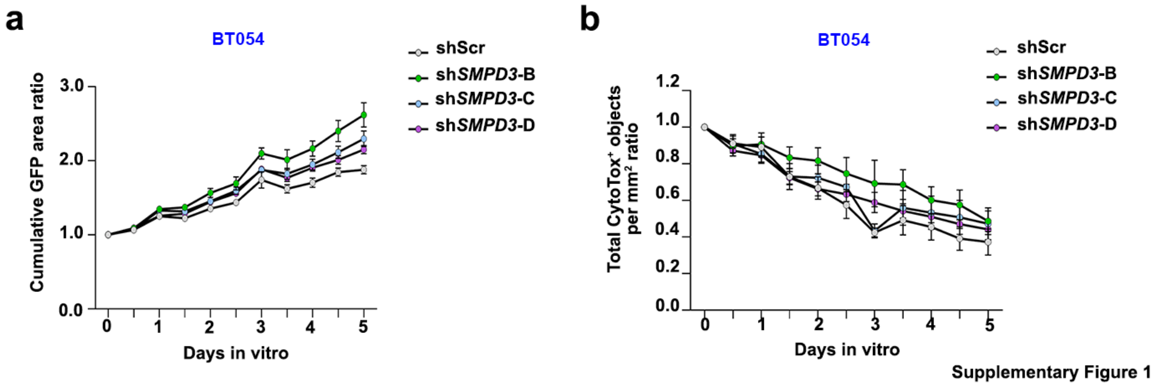

Supplementary Figure 1

**Supplementary Figure 2. BT054 IDH-O tumor cells are incapable of invading and proliferating in cerebral organoids (CO). a** Schematic representing the experimental setup to co-culture zSGreen-labeled BT054 cells with 10-month-old COs. **b-d** zSGreen-labeled BT054-CO coculture sections immunolabeled with zSGreen (green), MAP2 (red, **b**), SOX2 (red, **c**), and extracellular matrix markers COL4 (red, **d**, top) and LAM (red, **d**, bottom). Blue is DAPI nuclear counterstain. Scale bars: 200 µm.

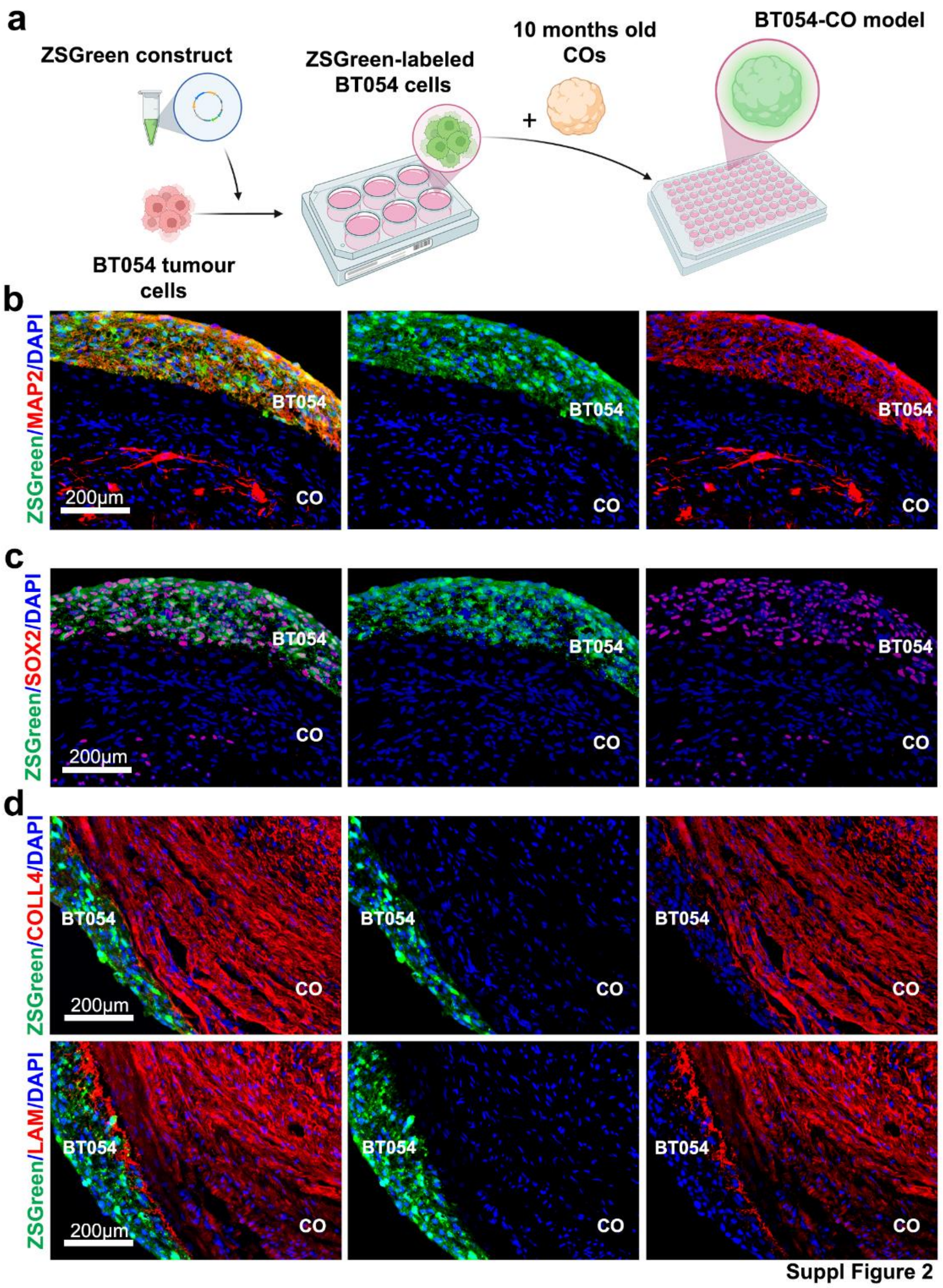

**Supplementary Figure 3. *Smpd3* knockdown reduces exosome secretion in BT088 IDH-O tumor cells. a** Western blot for whole HEK cell lysates co-transfected with shScr or sh*Smpd3* knockdown constructs. Three biological replicates of each sample set were loaded in varying amounts (5X/1X) to show knockdown efficiency. **b** BT088 cells transfected with shScr or sh*Smpd3* constructs were cultured for 10 DIV (BF, Brightfield). **c,d** Quantitation of GFP$^+$ tumoursphere numbers (**c**) and sphere diameter (μm) (**d**) after 10 DIV. Bars represent means ±s.e.m.. ***p<0.001.

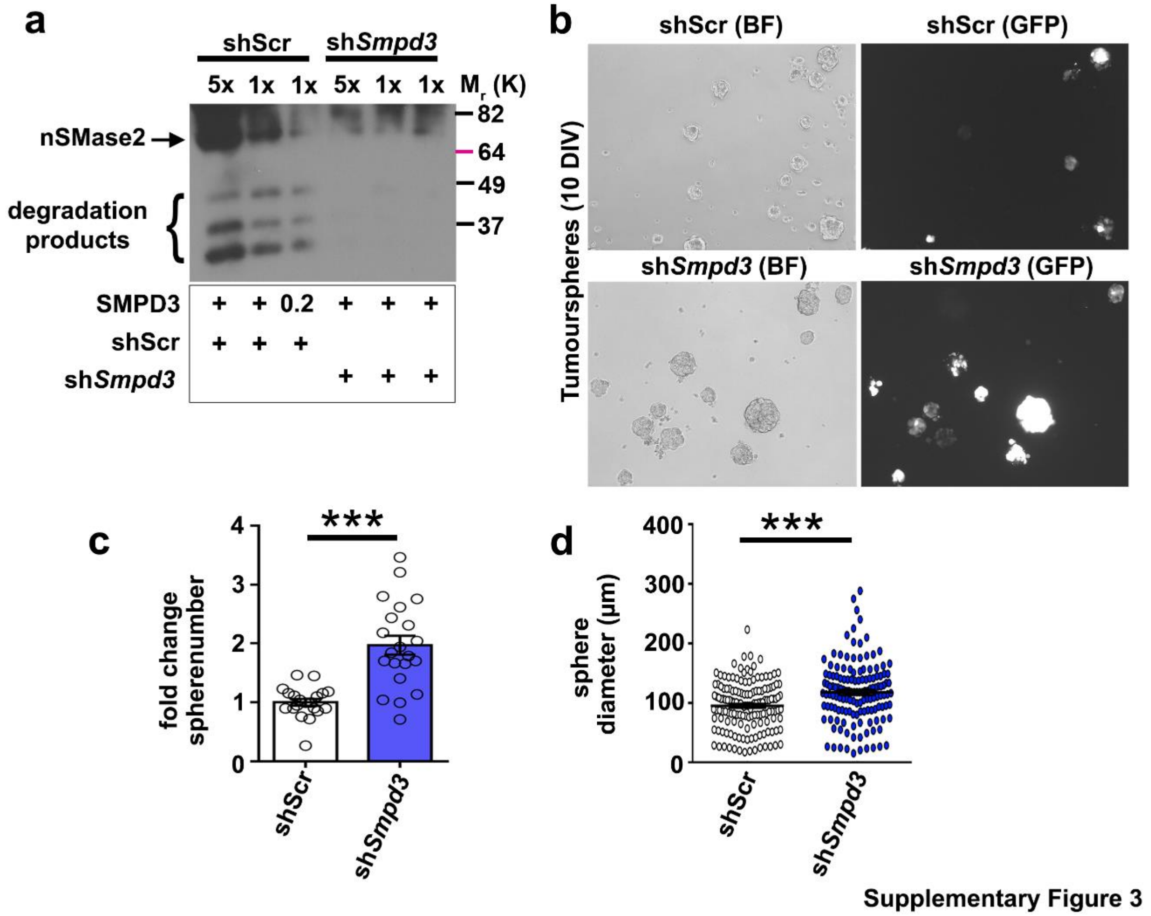

**Supplementary Figure 4. ODG cells secrete EVs mainly in the exosome size range. a** SEM of BT088 and BT054 cells. **b** Schematic of sequential ultracentrifugation protocol followed to isolate EVs from ODG cell condition media. **c** TEM of BT088 EVs. The red arrow marks an EV. A higher magnification image is shown to the right. **d** Nanosight tracking analysis of isolated BT054 and BT088 EVs. **e** CD9$^{+}$ EVs analyzed using nanoscale flow cytometry. **f** Western blots of BT088 whole-cell lysates (CL) and EV lysates were analyzed for markers of EVs (ALIX, CD9, CETP, Flotillin1), endoplasmic reticulum (Calreticulin, Calnexin), Golgi body (GM130), mitochondria (VDAC), and peroxisomes (PEX5). **g** Density gradient ultracentrifugation of BT088 EVs, with 8 fractions analyzed by western blots for expression of EV (ALIX, CD9, CETP) and non-EV markers (Calnexin, VDAC). **h** Nanosight tracking analysis of fraction 4 EVs. Scale bars: 2 µm in **a** (left) and 1 µm in **a** (right); 500 nm in **c** (left); 50 nm in **c** (right).

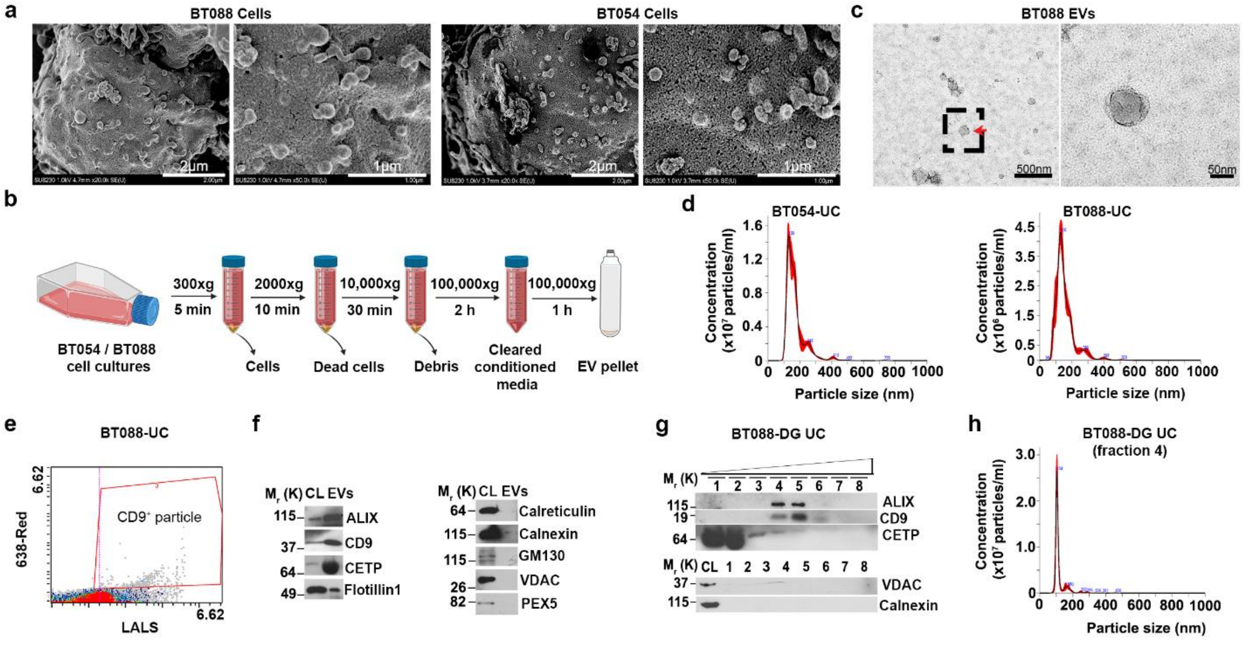


Suppl Figure 4

**Supplementary Figure 5. IDH-O malignant cells are mostly of oligodendrocytic lineage. a** IDH-O tumor resections (n=4) immunolabeled for IDHm (green) and oligodendrocyte marker (OLIG2, red). Blue is DAPI nuclear counterstain. **b** Percent ($OLIG2^+$) $IDHm^+$ or $IDHm^-$ cells normalized to total $OLIG2^+$ cells. Closed arrows represent cells positive for all markers, and open arrows show a lack of colocalization. Bars represent means ± s.e.m.. ****$p<0.0001$. Scale bars: 400 μm.

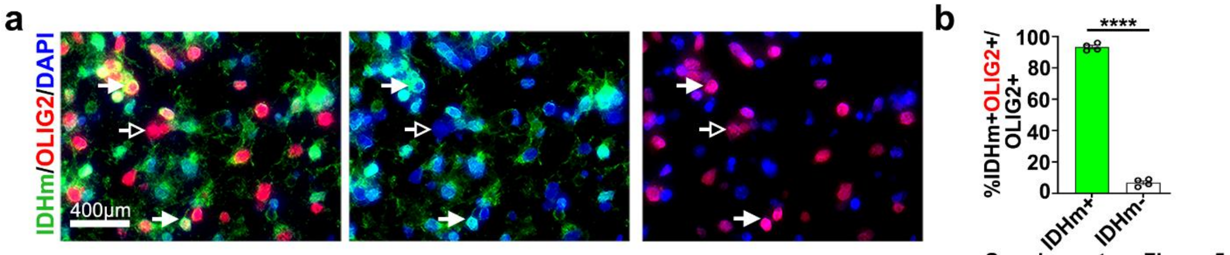

Supplementary Figure 5

**Supplementary Figure 6. Ectopic proliferation and cell death (apoptosis) of neoplastic and non-neoplastic cells in glioma subtypes' TME. a-h** IDH-mutant astrocytoma (**a, e**), IDH-mutant GBM (**b, f**), and IDH-wildtype GBM (**c, g**) tumor resections immunolabeled for IDHm (green) and KI67 (red, **a-c**) or CC3 (red, **e-g**). Blue is DAPI nuclear counterstain. **d** Comparison of percent proliferative $KI67^+$ cells in each tumor tissue normalized to the total number of $DAPI^+$ nuclei. **h** Comparison of percent apoptotic ($CC3^+$) cells in each tumor tissue normalized to the total number of $DAPI^+$ nuclei. Bars represent mean ± s.e.m.. *$*p<0.05$, $****p<0.0001$*. Scale bars: 400 μm.

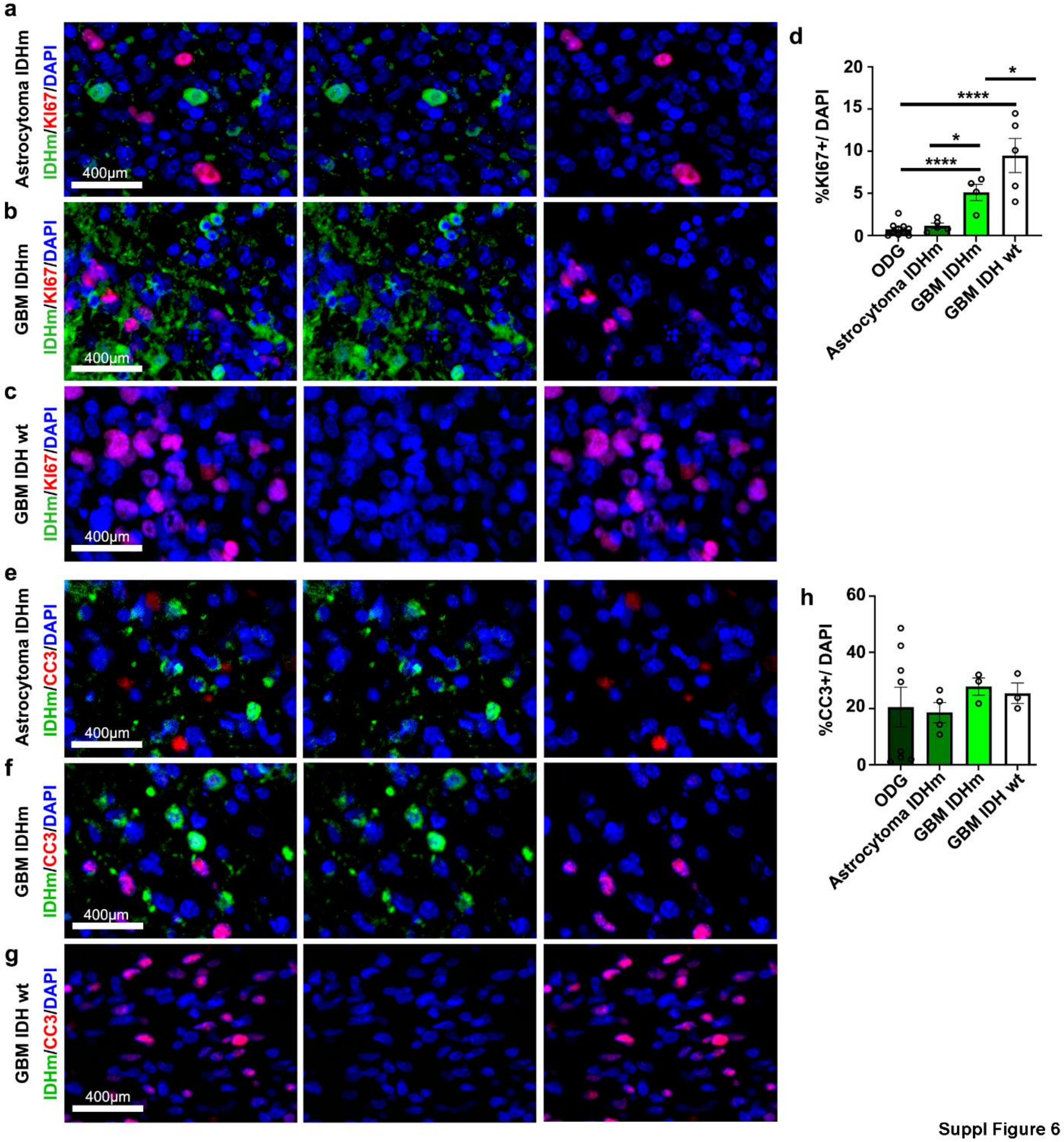


Suppl Figure 6

**Supplementary Figure 7. IDH-A-associated oligodendrocytes are enriched in ribosomal gene expression. a** tSNE plot of scRNA-seq data from IDH-A [1], showing cluster annotation, using CNVs to identify malignant cells, and cell type-specific markers to identify non malignant cells. **b-d** tSNE plots on which *MOG* (**b**) and *SEPP1* (**d**) expression were mapped. **c** GSEA of up-regulated genes in IDH-A-associated oligodendrocytes. (**e-h**) Comparisons of gene expression in IDH-A-associated oligodendrocytes and 'normal' brain oligodendrocytes.

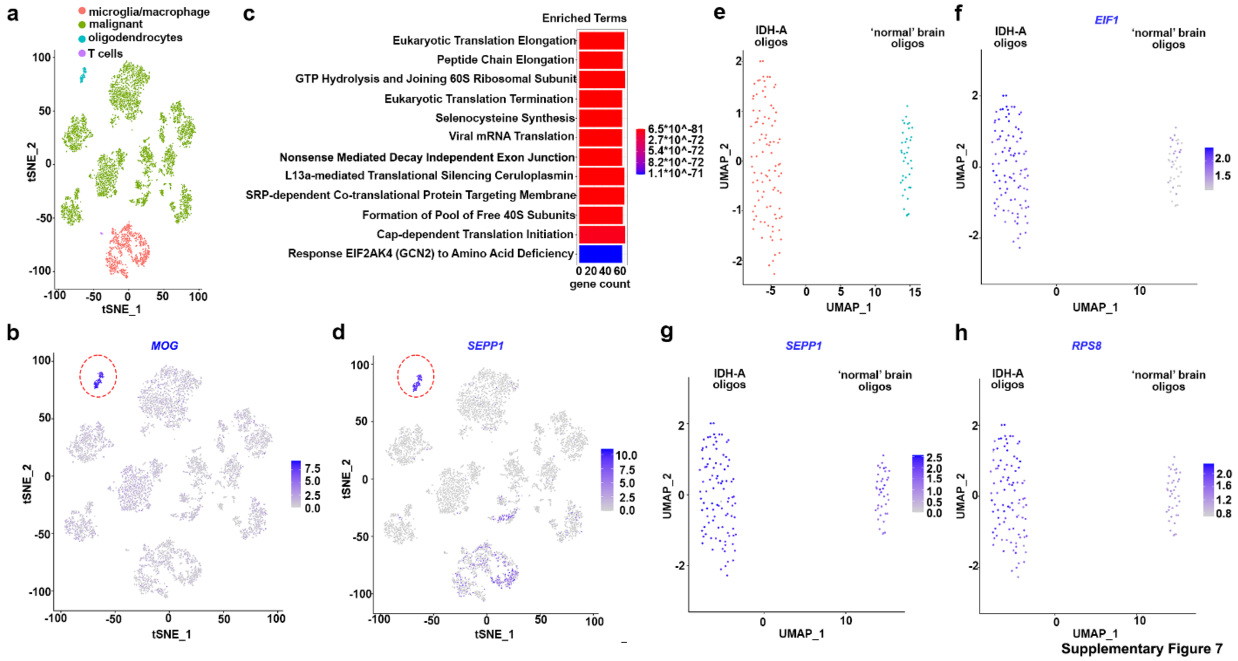


1 Venteicher, A. S. *et al.* Decoupling genetics, lineages, and microenvironment in IDH-mutant gliomas by single-cell RNA-seq. *Science* **355**, doi:10.1126/science.aai8478 (2017).
